# Visualising the cytoskeletal machinery in neuronal growth cones using cryo-electron tomography

**DOI:** 10.1101/2021.08.06.455451

**Authors:** Joseph Atherton, Melissa Stouffer, Fiona Francis, Carolyn A. Moores

## Abstract

Neurons navigate long distances and extend axons to form the complex circuitry of the mature brain. This depends on the coordinated response and continuous remodelling of the microtubule and F-actin networks in the axonal growth cone. Growth cone architecture remains poorly understood at nanoscales. We therefore investigated mouse hippocampal neuron growth cones using cryo-electron tomography to directly visualise their three-dimensional subcellular architecture with molecular detail. Our data show the hexagonal arrays of actin bundles that form filopodia penetrate the growth cone interior and terminate in the transition zone. We directly observe the modulation of these and other growth cone actin bundles by alteration of individual F-actin helical structures. Blunt-ended microtubules predominate in the growth cone, frequently contain lumenal particles and carry lattice defects. Investigation of the effect of absence of doublecortin, a neurodevelopmental cytoskeleton regulator, on growth cone cytoskeleton shows no major anomalies in overall growth cone organisation or in F-actin subpopulations. However, our data suggest that microtubules sustain more structural defects, highlighting the importance of microtubule integrity during growth cone migration.

**Summary statement:** Cryo-electron tomographic reconstruction of neuronal growth cone subdomains reveals distinctive F-actin and microtubule cytoskeleton architectures and modulation at molecular detail.

## INTRODUCTION

During development, neurons first form primary extensions that direct cell migration, and then extend axons to form synapses. These pathfinding processes are directed by the axonal growth cone in response to guidance cues and driven by an integrated network of microtubule (MT) and actin filaments (F-actin). Growth cone malfunction causes aberrant axonal pathfinding and incorrect wiring of the nervous system, and is the basis of a number of neurodevelopmental disorders including a range of intellectual disability conditions, autism and schizophrenia (Bakos et al., 2015; Kim and Kim, 2020; Stoeckli, 2018). For example, mutations in the protein doublecortin (DCX in humans, Dcx in mouse), which is a regulator of both growth cone F-actin and MT organisation (Deuel et al., 2006; Friocourt et al., 2003; Fu et al., 2013; Jean et al., 2012; Tint et al., 2009), cause neurodevelopmental lissencephaly (Wynshaw-Boris et al., 2010). Understanding the organisation and regulation of the neuronal growth cone cytoskeleton can thus shed light on the molecular basis of numerous diseases (Roig-Puiggros et al., 2020).

Distinct growth cone regions have been characterised based on their appearance, organelle distribution and cytoskeletal composition (Sup. Fig. 1A, B). The axon shaft that precedes the growth cone is filled with parallel, tightly bundled MTs - large organelles such as mitochondria are common and F-actin is relatively sparse (Dent et al., 2011). The axon connects with the central domain (C-domain), which includes the so-called axon wrist, and, while also containing large organelles and MTs, C-domain MTs are more dispersed and dynamic (Blanquie and Bradke, 2018). The front region of the growth cone - the peripheral domain (P-domain) - is flat and organelle-free. It is constituted of F-actin bundle-containing protrusions called filopodia, together with the branched F-actin network containing lamellipodium. The P-domain is separated from the C-domain by the transition zone (T-zone) which, as its name suggests, is a region of transition between the F-actin dominated outer regions of the growth cone and the MT-rich axon-proximal regions. Contractile Factin arcs formed by non-muscle myosin II run parallel to the front of the growth cone in the T-zone and help drive cellular motility (Burnette et al., 2008; Burnette et al., 2007; Schaefer et al., 2002; Zhang et al., 2003). MTs only rarely grow from the T-zone into the P-domain, and are often guided by F-actin bundles (Gordon-Weeks, 1991; Sabry et al., 1991; Schaefer et al., 2002; Schaefer et al., 2008). Furthermore, MTs that are found in the periphery are dynamic and unstable - exploratory MTs that are linked to F-actin retrograde flow are often induced to buckle and break in the T-zone (Schaefer et al., 2002; Schaefer et al., 2008; Waterman-Storer and Salmon, 1997).

There are a number of open questions concerning the molecular basis for growth cone cytoskeleton regulation and coordination. The precise organisation of non-filopodia P-domain F-actin and its connectivity and communication with the T-zone is not well understood. In particular, the mechanism by which filopodial F-actin bundles are disassembled, and how actin arcs are generated in the T-zone is not clear. In addition, it is not known whether different growth cone MT populations have distinct structural signatures, and, although the involvement of a number of MT-actin crosslinkers have been proposed (Geraldo and Gordon-Weeks, 2009; Lowery and Van Vactor, 2009), how peripheral MTs in growth cones associate with or are influenced by different arrays of F-actin is currently unclear. Nanoscale insights into neuronal growth cone organisation can provide key insights to address some of these outstanding questions.

Cryo-electron tomography (cryo-ET) provides a window into the growth cone interior by enabling visualisation of intact frozen-hydrated cellular samples; threedimensional (3D) reconstruction enables ultrastructural characterisation of molecular components *in situ* (Turk and Baumeister, 2020). We therefore used cryo-ET and machine learning-based segmentation and visualisation to investigate the complex molecular organisation of sub-compartments within mouse hippocampal neuron growth cones and the part of the axon that leads in to them. We focused particularly on their underlying cytoskeletal architectures vital to axon pathfinding. We also compared the growth cone ultrastructure in mouse wild-type neurons with those of Dcx knockout neurons, to evaluate the effect of Dcx loss on the cytoskeleton.

## RESULTS

### Cryo-electron tomography of primary hippocampal neuron growth cones

We chose mouse primary hippocampal neurons as a model system because the biology of their growth cone formation and function is well studied, and because the Dcx knockout phenotype in mice manifests itself most prominently in a hippocampal malformation (Corbo et al., 2002; Kappeler et al., 2007). After 2-3 days of *in vitro* culture on laminin/poly-D-lysine, these cells exhibit several short minor dendritic processes, together with an extended axon, tipped by a growth cone (Sup. Fig. 1A; reviewed in (Banker, 2018). Dcx mostly colocalises with MTs towards the outer ends of processes and in the growth cone C-domain (Bielas et al., 2007), but less intense Dcx staining is also seen in the P-domain colocalised with actin (Sup. Fig. 1A,B) (Fu et al., 2013; Tint et al., 2009).

Mouse primary hippocampal neurons were therefore grown on laminin/poly-D-lysine-treated holey carbon electron microscopy (EM) grids for 2-3 days. Grids were vitrified, growth cones identified at low magnification (Sup. Fig. 1C), tilt series collected and tomographic volumes were calculated, allowing clear visualisation of the interior of the neuron (Sup. Fig. 1D, Sup. Video 1).

### Visualisation of the axonal growth cone molecular landscape

We used neural network-based approaches to perform semi-automated segmentation of the growth cone tomograms, enabling 3D visualisation of their molecular organisation (Sup. Fig. 2). We also used the segmentation analysis to quantitatively assess the molecular distribution of cellular components within the different regions of the growth cone (Fig. 1), consistent overall with previous observations (Geraldo and Gordon-Weeks, 2009). There is a sharp drop of MT density from the C-domain through the T-zone into the P-domain, and this is the inverse of F-actin distribution, where diverse F-actin arrays dominate the growth cone periphery (Fig. 1A). Larger membrane-bound organelles are relatively common in the C-domain (Fig. 1B), including ER networks (Fig. 1C), large vesicles (>100 nm diameter, Fig. 1D) and components of the endolysosomal system (Fig. 1E). In the P-domain, these cellular components are very rare and are perhaps excluded because of their physical dimensions. The distribution of smaller (<100 nm diameter) vesicles is more variable among different growth cones (Fig. 1F). While a variety of small vesicles were observed in the P-domain and even in filopodia, those with clathrin-coats were only observed in the C-domain (Fig. 1G). Individual ribosome and polysome distributions remain relatively constant throughout the growth cone, except in the filopodia where they are more abundant, especially compared to the rest of the P-domain (Fig. 1H).

**Figure 1.**
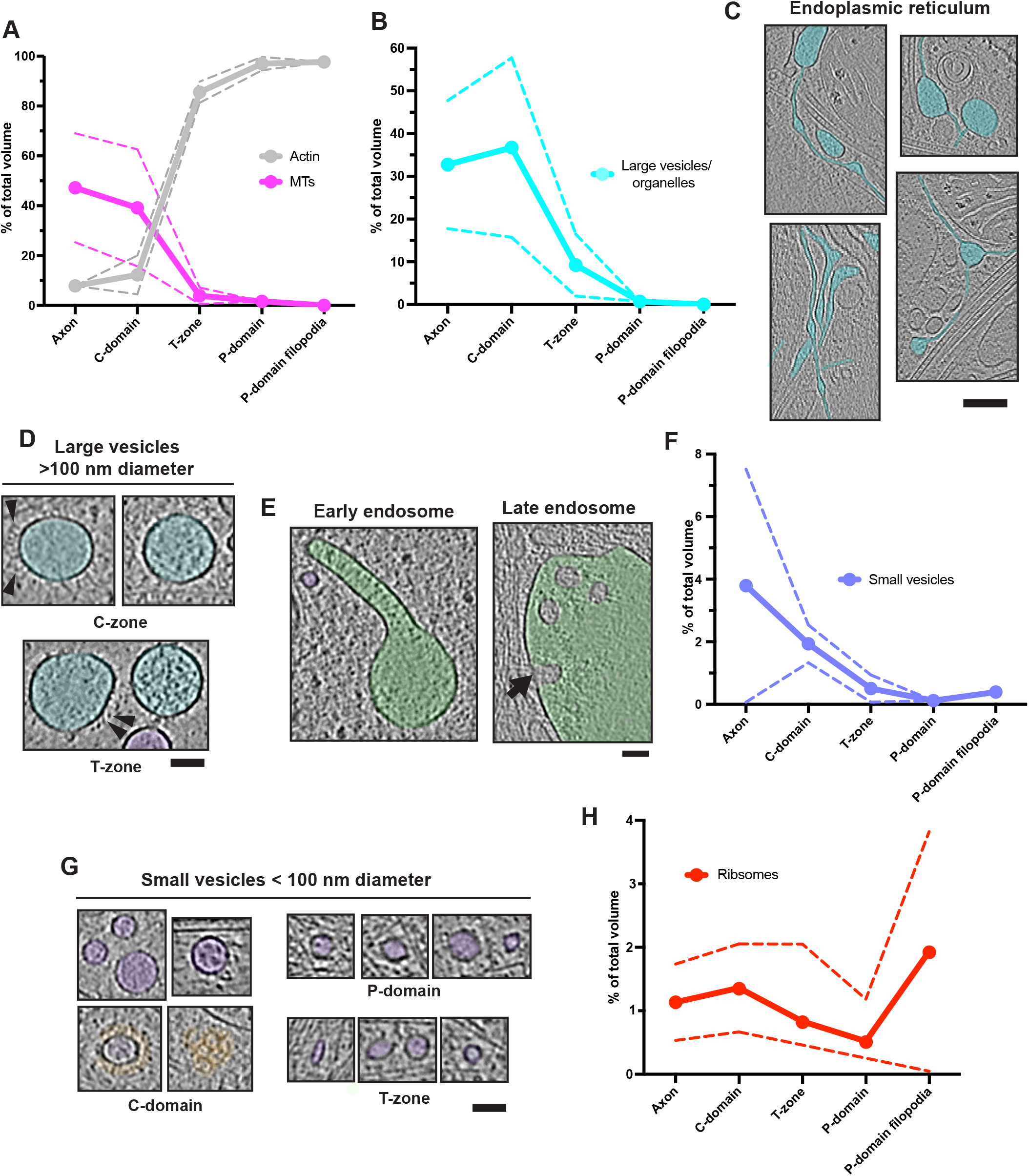
Quantitative analysis of growth cone component distribution. **A)** Growth cone distribution of segmentation volumes of MTs and F-actin. **B)** Segmentation volume of large vesicles and organelles (including mitochondria). **C)** Smooth ER in the C-domain, false coloured in cyan. **D)** Large vesicles, false coloured cyan. Arrowheads, embedded proteins. **E)** Endolysosomal components in the C-domain (left, early/sorting endosome, right, late endosome/multi-vesicular body), false coloured in green. Arrow indicates inward budding site. **F)** Segmentation volume of large vesicles and organelles. **G)** Small vesicles <100 nm diameter, false coloured in purple. Exemplar clathrin coats are false coloured in orange. **H)** Segmentation volume of ribosomes. Panels C-E and G show longitudinal views in tomograms (4x binned). Absolute segmentation volumes were measured in Chimera (Pettersen et al., 2004), and relative segmentation volume for each feature indicated is expressed as a % of the total volume of all these features combined. Axon, n = 2, C-domain, n = 3, T-zone, n = 3, P-domain, n = 2, P-domain filopodia, n = 2, dashed lines indicate S.D. Scale bars: C = 100 nm; D = 60 nm; E = 60 nm; G = 60 nm.

### Pseudo-crystalline organisation of neuronal filopodia actin bundles

The molecular detail visible in our tomograms allowed us to characterise the diverse manifestations of the actin cytoskeleton that dominate outer growth cone regions. At the very front of the growth cones, several filopodial regions are typically captured in each tomogram (Fig. 2A, Sup. Fig. 2D), the dominant feature of which are tight bundles of F-actin filaments (Fig. 2B). Surrounding the bundles, individual actin filaments lie beneath the membrane, some running roughly perpendicular to the filopodial axis (Fig 2A,B), and some which incorporate into the bundles. Smaller filopodial extensions (< 400 nm diameter) contain a single F-actin bundle, while larger filopodia contain 2-3 discrete bundles which exchange filaments or coalesce (Fig. 2B). Other non-cortical, non-bundled actin is also found flanked by bundles, or between bundles and the membrane (Fig. 2B).

**Figure 2.**
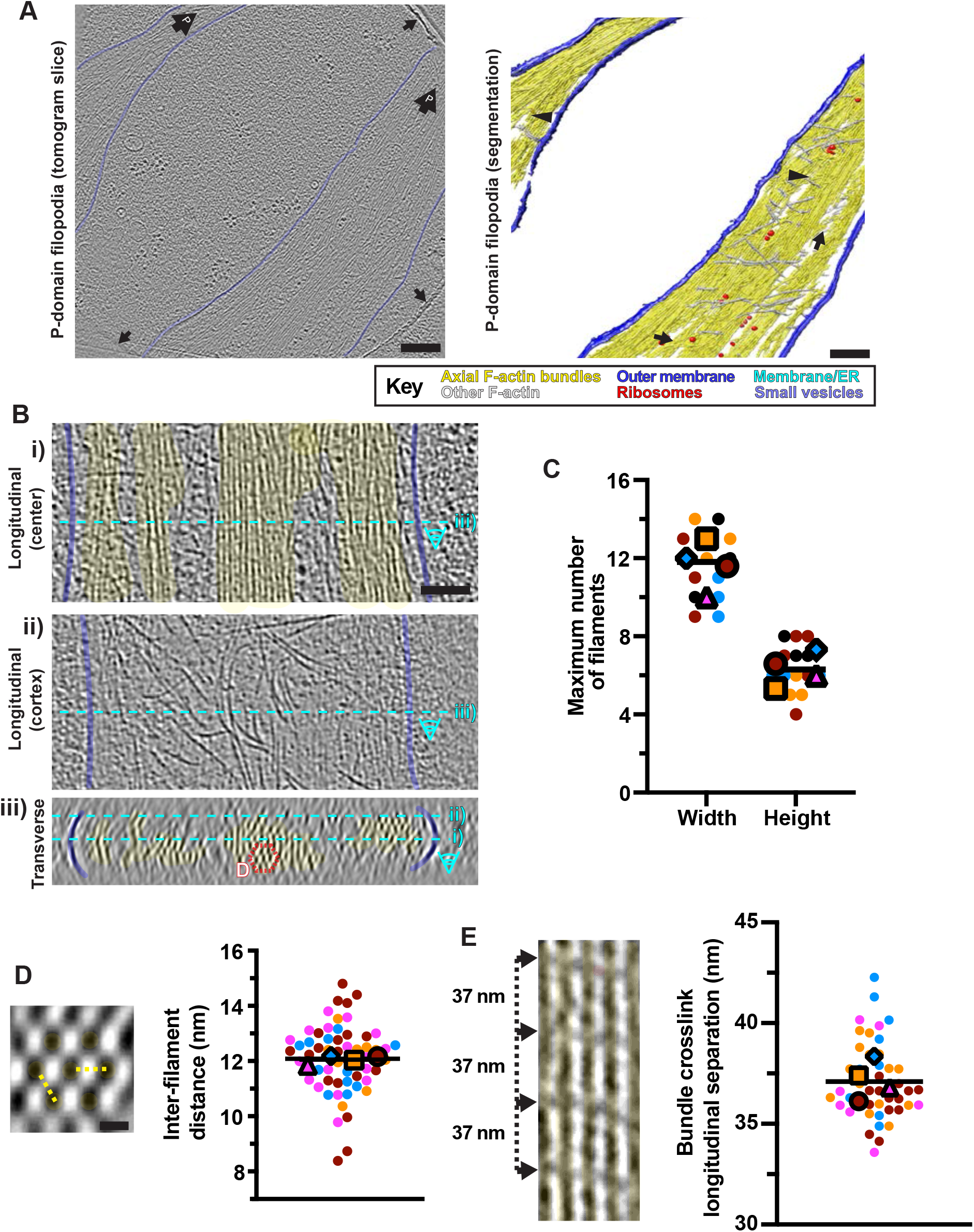
Ultrastructure of filopodial F-actin arrays. **(A)** Central tomogram section (4x binned), left, segmentation, right, of 2 filopodia. Left, large black arrow containing a ‘P,’ position of cell front; black arrows, carbon hole edges; right, black arrows, F-actin filaments connecting bundles; black arrowheads, indicate merging of F-actin bundles. Key indicates false colouring scheme. **(B)** Tomogram sections (4x binned) of a large filopodium showing the (i) central region, (ii) cortex and (iii) cross-section (~20 nm depth). Cyan dashed lines indicate how panels relate. Dashed red hexagon indicates a hexagonal F-actin bundle unit (panel D). **(C)** Super-plot (Lord et al., 2020) of maximum width/height of filopodial bundles. Each data point represents a filopodial bundle (N=14 from 4 tomograms in different colours); line = overall median. Mean tomogram values shown in respective colours with different larger shapes. Overall mean width, 11.6 ± 1.7 S.D, mean height 6.4 ± 1.2 S.D filaments. **(D)** Left; transverse view through a filopodial F-actin bundle (~20 nm depth), showing its characteristic hexagonal arrangement. Yellow dashed lines, exemplar inter-filament distances. Right, Superplot of inter-filament distances in filopodial bundles. Each data point represents a separate adjacent filament pair (N=60 from 4 tomograms, shown as different colours). Mean values for each tomogram are shown in their respective colours with different larger shapes. Line indicates the overall mean (12.1 nm ± 1.2 S.D). **(E)** Left, a single cross-linked sheet in a filopodial bundle (4x binned), showing ~37 nm crosslink separation, indicated with arrows; Right, Super-plot of longitudinal crosslink separation in P-domain F-actin bundles. Each data point represents an individual pair of cross-links along the F-actin longitudinal axis (N=44 from 4 tomograms, shown as different colours). Mean values for each tomogram are shown in their respective colours with different larger shapes. Line indicates the overall mean (37.1 nm ± 1.9 S.D). Scale bars: A = 200 nm, B = 100 nm, D = 10 nm.

Filaments within all the filopodia we observed are arranged in a characteristic 2D hexagonal, pseudo-crystalline array (Fig. 2B) (Kureishy et al., 2002), although this arrangement is not strictly maintained - the positions of individual filaments meander within the larger bundles, and bundle thickness varies (Sup. Video 2). We observed bundles of 4-8 filaments in height and 9-14 filaments in width (Fig. 2C), with an inter-filament spacing of ~12 nm (Fig. 2D), and connecting density between adjacent bundle filaments was spaced at ~37 nm intervals along the diagonals of the hexagonal array (Fig. 2E).

This characteristic hexagonal filament organisation acts as a molecular signature to track filopodia-derived bundles into the P-domain cytoplasm. Our data show how they penetrate and integrate into the complex actin-rich environment of the P-domain (Fig. 3A, Sup. Fig. 2C). The filament arrangement is very consistent between different bundles, both within the same filopodia, between different filopodia and in bundles that extend into the P-domain (Sup. Fig. 3). To better understand bundle molecular organisation, we performed sub-tomogram averaging of a 7-filament hexagonal repeating unit (Fig. 3B-D). In the resulting 3D reconstruction (nominal resolution 29 Å, Sup. Fig. 4), individual 5.5 nm actin subunits are distinguishable, and the filaments exhibit a half helical pitch distance of ~37 nm (Fig. 3C). Crosslinking densities are regularly spaced and connect the central filament (filament 0, Fig. 3B) with four of the six surrounding filaments (filaments 2,3,5,6, Fig. 3B-D; Sup. Video 3). Density also connects the peripheral filaments 1-2, 3-4, 4-5 and 6-1 (Fig. 3B). The distribution of these cross-links relates to the helical structure of the individual filaments. When viewed as a 2D slice, cross-links join filament 0 to filaments 2 and 5 every 37 nm and, orthogonally, filament 0 to filaments 3 and 6 (Fig. 3C,D). Cross-links only occur where the outer edge of adjacent, helically aligned filaments approach other (Fig. 2E, 3C), and therefore the two cross-linked orthogonal planes (2-0-5, 3-0-6) do not have their cross-links at the same position along the filament axis (Fig. 3D). Because there are no cross-links in the 1-0-4 plane, there is a repeating pattern of cross-links along the bundle axis, joining the central filament (0) to filaments 3 and 6 followed by a 12.3 nm gap, then connections to filaments 2 and 5, followed by a 24.6 nm gap to the next cross-links with filaments 3 and 6 (Fig. 3D, Sup. Video 3). This pattern produces an approximate stoichiometry of 1 cross-linking density per ~5 actin monomers.

**Figure 3.**
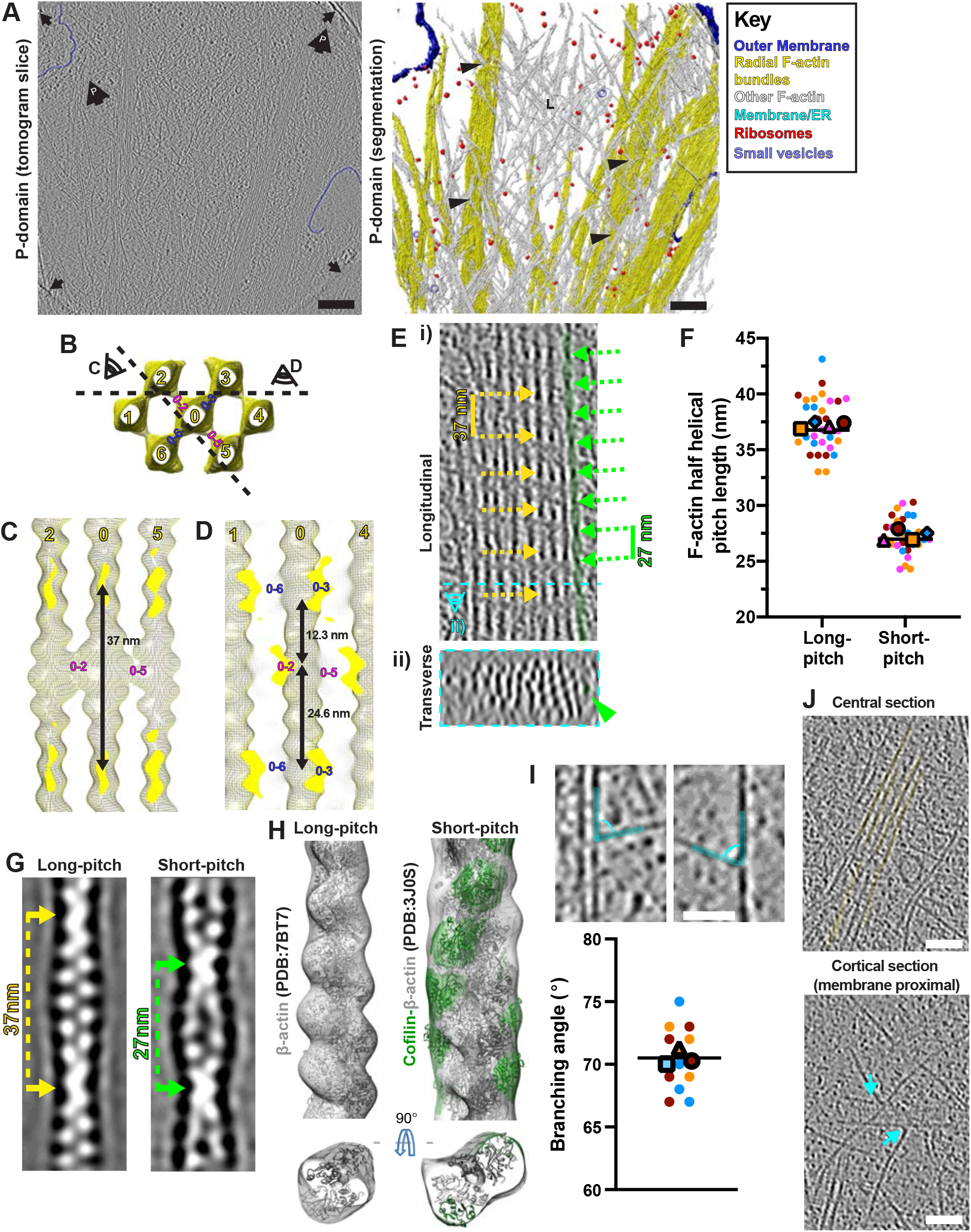
Complexity and modulation of F-actin arrays in growth cone P-domains. **(A)** Central tomogram section (4x binned), left, semi-automated segmented rendering (right) of a P-domain region. Left, large black arrow containing a ‘P,’ position of cell front; black arrows, carbon hole edges; right, L= Lamellipodial; black arrowheads, branched F-actin networks surrounding F-actin bundles. Key indicates false colouring scheme. **(B-D)** Subtomogram average (yellow mesh) of hexagonal P-domain bundles. **(B)** transverse section with filaments numbered to aid description and 90° rotated viewpoints shown in C and D indicated. Individual cross-links are labelled according to numbered filament they connect. **(C)** 37 nm separation of cross-links along single 2D plane connecting 2-0-5 filaments. **(D)** Cross-linking superstructure showing cross-links connecting 2-0-5 and 3-0-6 filaments. Individual cross-links are labelled as in panel B. **(E**) i) Longitudinal, ii) transverse (~20 nm depth) views of a large F-actin bundle in a 2x binned P-domain tomogram. Dashed cyan line in i) illustrates position of transverse section in ii). Canonical ~37 nm Factin half helical pitch lengths indicated with yellow dashed arrows, shorter, rarer ~27 nm F-actin half helical pitch lengths indicated with green dashed arrows. A short-pitch filament length is false coloured in green and additionally indicated in ii) with a green arrowhead. **(F)** Super-plot of P-domain F-actin longitudinal half helical pitch lengths. Individual points indicate single measurements along a filament axis, colours indicate different filaments. Mean values for each filament are shown in their respective colours with different larger shapes. Individual lines indicate overall median values. Long-pitch filaments; mean = 37.2 nm ± 2.4 S.D, N=32, 4 filaments (3 tomograms); short-pitch filaments; mean = 27.3 nm ± 1.6 S.D, N=30, 4 filaments (3 tomograms). **(G)** Projections through subtomogram averages of long-pitch (left) and short-pitch (right) filaments showing filament half pitch distances. Long-pitch filament = average of 1461 volumes. Short-pitch filament = average of 621 volumes. Volumes were low-pass filtered to their estimated resolutions at the Fourier Shell Correlation (FSC) 0.5 criterion (Sup. Fig. 4). **(H)** Subtomogram averages of long-pitch filaments (left, mesh density) and short-pitch (right, mesh density) filaments with fitted F-actin (PDB: 7BT7) or F-actin-cofilin (PDB: 3JOS) models respectively. FActin and cofilin models are coloured in grey and green respectively with mesh density coloured to match the underlying models (within 10 Å). **(I)** P-domain tomogram sections (4x binned) illustrating F-actin branching from bundles (top left) and single filaments (bottom left), indicated with cyan false colouring. Right; Superplot of P-domain branching angles. Each data point represents an individual branching structure (N=12 from 3 tomograms, shown as different colours). Mean values for each tomogram are shown in their respective colours with different larger shapes. Line indicates the overall mean (70.1° ± 2.6 S.D). **(J)** Longitudinal sections of a 2 x binned P-domain tomogram through left, a central region showing radial Factin bundles in false yellow colour and right, the corresponding overlying cortical region, showing a F-actin branching points at the cell cortex. Branching points are indicated with cyan arrows. Scale bars: A = 200 nm, I = 40 nm, J = 50 nm.

### Modulation of F-actin arrays in growth cone P-domains

Strikingly, despite the conservation of the core structure of the filopodial bundles, a minority of filaments - either within the bundles, particularly at their periphery, or in separate filaments in the P-domain (Fig. 3E, Sup. Fig. 5A-C) - have a distinct appearance and a shorter half helical pitch length of ~27 nm (Fig. 3E,F). There is no evidence of filaments with intermediate half helical pitch length (Fig. 3F), and the ~27 nm half pitch filaments tend to be shorter and lack regular cross-links to adjacent filaments (Fig. 3E, Sup. Fig. 5). Furthermore, these short-pitch filaments generally meander more within the bundle, and often appear to be disrupting the regularity of, the bundle with which they are associated (Sup. Fig. 5A,B). Some individual filaments clearly transition from one helical structure to the other (Sup. Fig. 5C).

Sub-tomogram averages of the ~37 nm and ~27 nm half helical pitch filament populations (Fig. 3G & Sup. Fig. 4) recapitulate the actin twists and helical pitches visible in 2D slices (Fig. 3G). Further, the ~27nm half helical pitch filament reconstruction suggests that additional protein(s) is/are bound. Based on extensive prior knowledge of the role of cofilin/ADF in actin cytoskeleton regulation (Gungabissoon and Bamburg, 2003) and the known structural twist imposed by cofilin/ADF on F-actin filaments (Galkin et al., 2011; Galkin et al., 2001; Huehn et al., 2020; McGough et al., 1997; Tanaka et al., 2018), we rigid-body fitted F-actin-cofilin (PDB: 3JOS (Galkin et al., 2011) within this density (Fig. 3H, Sup. Video 4), which exhibited a very good match (0.87 model to map cross-correlation calculated in Chimera (Pettersen et al., 2004)). The known binding site of cofilin - longitudinally between actin subunits along F-actin - corresponded well with the non-actin density in the reconstruction. Our ability to visualise this density suggests high binding protein occupancy which, together with the observation of extended stretches of ~27nm half helical pitch in the raw tomograms, is consistent with the reported cooperativity of cofilin binding *in vitro* (Huehn et al., 2018).

The P-domain also contains branched F-actin networks formed from a mixture of single filaments and small bundles between the filopodia-derived actin bundles (Fig. 3A). Branching originates from the sides of single filaments and, less frequently, from bundles, all at ~70° angles (Fig. 3I), consistent with Arp2/3 complex activity. While this branched network is distributed throughout the P-domain, the branch points themselves are primarily located close to the dorsal and ventral membranes (Fig. 3J).

### The mixed cytoskeletal economy of the T-zone

The T-zone cytoskeleton is characterised by 1) the termination of radially organised P-domain-derived F-actin bundles and 2) the termination of the majority of MTs originating from the C-domain. Filopodia-like F-actin bundles extend into the T-zone (Fig. 4A), oriented radially towards the front of the growth cone. Although these bundles containing fewer filaments (maximum height of ~4 filaments, maximum width of 8 filaments), they exhibit the characteristic average inter-filament spacing, roughly hexagonal distribution, and ~37nm half helical pitch lengths seen in the P-domain (Fig. 4B, C). These bundles tend to taper within the T-zone, with subbundles and single filaments often bending off perpendicularly to the main bundle axis (Fig. 4A, D). However, even when filaments are only cross-linked in 1D, they are regularly spaced and connected by crosslinking density at ~37nm intervals (Fig. 4D,E). As was also observed in the P-domain, a minority of filaments, typically shorter and at the periphery of the bundles, exhibit a ~27 nm half helical pitch length (Fig. 4F, Sup. Fig. 5D).

**Figure 4.**
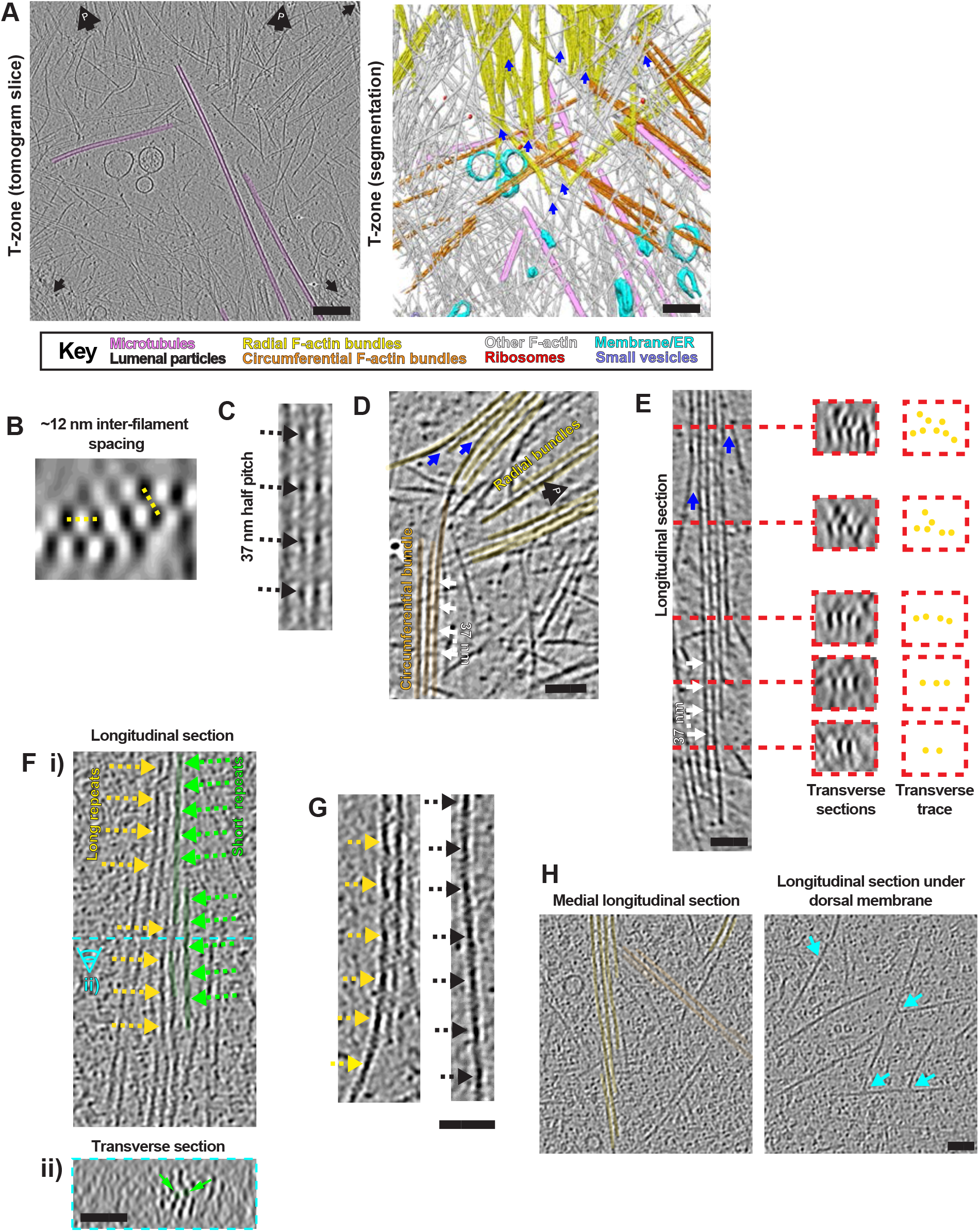
Diverse F-actin arrays in neuronal growth cone T-zone. **(A)** Central tomogram section (4x binned), left, semi-automated segmented rendering, right, of a T-zone region. Left, large black arrow containing a ‘P,’ position of cell front; black arrows, black arrows, carbon hole edges. Key indicates false colouring scheme. **(B)** Transverse section through a radial F-actin bundle in a T-zone tomogram (4x binned), showing hexagonal filament organisation; yellow dashed lines indicate ~12 nm inter-filament distances. **(C)** Longitudinal section through a radial F-actin hexagonal bundle in a T-zone tomogram (2x binned), showing 37 nm half helical pitch lengths indicated by arrows. **(D)** Longitudinal section through a tomogram (4x binned) of a radial F-actin bundle taper and a derived circumferential F-actin bundle, with white arrows indicating F-actin cross-links. White arrows indicate 37 nm inter-crosslink spacing along the filament axis. **(E)** Longitudinal (left) and transverse (centre, ~20 nm depth) sections, indicated with red dashed lines, of a tapering radial F-actin bundle in a T-zone tomogram (4x binned). A traced representation (right) illustrates bundle tapering. White arrows indicating Factin cross-links. White arrows indicate 37 nm inter-crosslink spacing. **(F)** i) Longitudinal and ii) transverse views of a tapering radial F-actin bundle in a T-zone tomogram (2x binned). In i) distal and proximal are at the top and bottom respectively. Dashed cyan line in i) indicates the position of transverse section in ii). ~37 nm and ~27 nm half helical pitch lengths are indicated with yellow and green dashed arrows, respectively. Short-pitch filament lengths are false-coloured in green and their transverse positions within the bundle indicated in ii) with a green arrowhead. **(G)** Longitudinal tomogram sections (2x binned) of a radial F-actin bundle taper (left) and an F-actin bundle-derived dislocated single filament (right), dashed arrows indicate long F-actin half helical pitch lengths. **(H)** Longitudinal sections of a T-zone tomogram (2x binned) through left, a central region showing radial and circumferential F-actin bundles in false yellow and orange colour respectively, right, the corresponding overlying cortical region, showing F-actin branching points at the cell cortex. Branching points are illustrated with cyan arrows. Blue arrows in A, D, E indicate points where filaments bend-off tapering radial Factin bundles. Scale bars: A = 200 nm, D-E = 50 nm, F-H = 40 nm.

Compared to the P-domain, the radial actin bundles are less dense in the T-zone, (Fig. 4B,C), and are interspersed among smaller bundles orientated parallel to the growth cone circumference (Fig 4A & Sup. Fig. 2B). Additionally, numerous individual filaments run parallel or perpendicular to the radial bundles (Fig. 4A, Sup. Fig. 2B). These distinct T-zone actin populations are often clearly derived from axial bundles (Fig. 4A,D-G), although their overall organisation is looser and apparently less precisely organised, particularly at the cell membrane. Circumferentially organised F-actin also runs roughly perpendicular to the periphery-oriented MTs, but does not appear to block their trajectory (Fig. 4A). Branched actin arrays consistent with Arp2/3 activity are also visible near the T-zone membrane (Fig. 4H). Thus, while tomograms of the growth cone T-zone show that the F-actin network is overall less dense than the P-domain, they also clearly demonstrate that the actin cytoskeleton is organised in distinct populations, mediated by the coordinated activities of regulatory proteins.

### Growth cone microtubule architecture and interactions with F-actin

MTs are most prevalent in the C-domain (Sup. Fig. 2A-C), and all observed MTs throughout the growth cone exhibit a canonical 13-protofilament architecture (N=141, 11 tomograms) (Fig. 5A). Using the handedness of MT transverse segment rotational averages (Bouchet-Marquis et al., 2007; Sosa and Chretien, 1998), we could also show that the vast majority of growth cone MTs were orientated with their plus end direction towards the periphery with only a few exceptions (Fig. 5B), as found previously in murine axons (Baas and Lin, 2011). A further subset of MTs in the T-zone and C-domain were either short, straight and perpendicular to the axonal axis (with both ends visible), or were long and looped ~180° such that they bent back on themselves, and thus neither of their ends pointed peripherally (Fig. 5B (N/A subset), Fig. 6A, Sup. Fig. 2A).

**Figure 5.**
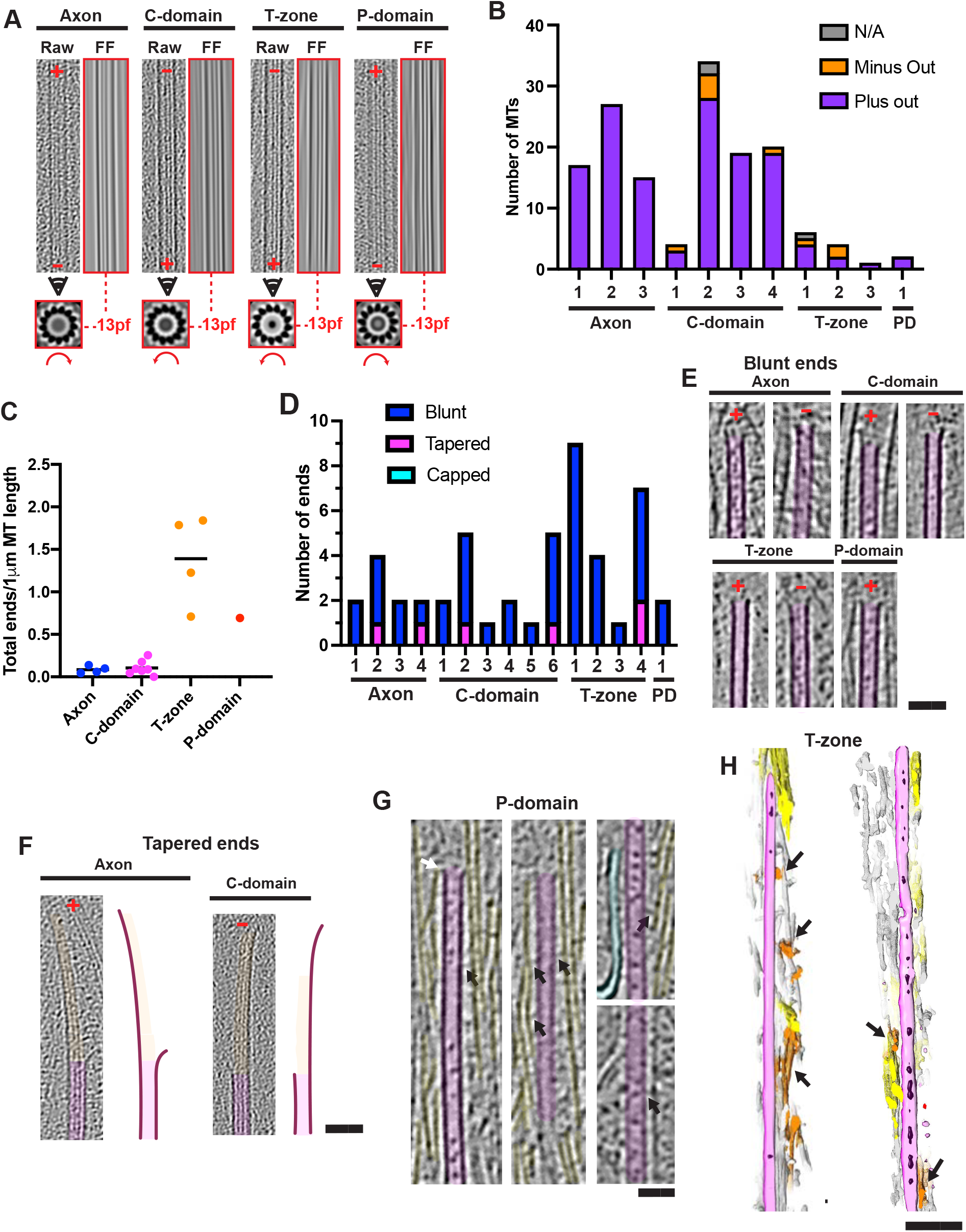
Organisation and architectures of microtubules within growth cones. **(A)** Exemplar MTs illustrating protofilament number and polarity. Each set of 3 images includes a ~30 nm thick longitudinal section through a MT volume (left, 2x binned) with its corresponding image Fourier filtered at the origin (right, red box) to show 13-protofilament moiré patterns. Below is a corresponding MT rotational average of a 30 nm thick section viewed towards the cell periphery, showing protofilament number and handedness (curved red arrow). When viewed from the minus end or plus end, rotational average images exhibit clockwise or anticlockwise slew respectively. In longitudinal sections the growth cone periphery is towards the top of the images, plus and minus end directions are indicated with a red ‘+’ and ‘−’ respectively, and the consensus protofilament (pf) architecture is indicated between dashed red lines. **(B)** MT polarity relative to neuron periphery was quantified in individual tomograms. MTs assigned ‘N/A’ were either perpendicular to the axon axis or were bent ~180°. Axon N=59 total plus-end peripheral MTs, 0 minus-end peripheral from 3 tomograms (each a different cell); C-domain N=69 total plus-end peripheral MTs, 6 minus-end peripheral, 2 n/a from 4 tomograms (each a different cell); T-zone N=7 total plus-end peripheral MTs, 3 minus-end peripheral, 1 n/a from 3 tomograms (each a different cell), P-domain (PD) N=2 total plus-end peripheral MTs, 0 minus-end peripheral, 0 n/a from 1 tomogram. **(C)** Frequency of MT ends per 1 μm MT length in individual tomograms. Each data point represents a separate tomogram; axon=4, C-domain=7, T-zone=4, P-domain=1. Line indicates the mean from all tomograms for each region. 50 MT ends were found in a total of 318 μm MT length. **(D)** Quantification of the absolute number of blunt and tapered ends in individual tomograms (number of tomograms of axon=4, C-domain=7 (one tomogram contained no ends), T-zone=4, P-domain=1). **(E)** Longitudinal views of blunt and tapered **(F)** MT ends. Blunt ends - 10 nm thick slices through 4x binned tomograms; tapered ends - 30 nm thick slices through 2x binned tomograms, false coloured in magenta and orange respectively. A traced representation of each tapered MT end is shown to its right. Plus and minus ends are indicated on images with a red ‘+’ and ‘−’. **(G)** Four longitudinal sections through regions of a single MT associated in parallel with P-domain F-actin bundles. Black and white arrows indicate cross-links between the MT shaft or MT tip and F-actin bundles respectively, and cyan indicates membranous ER. **(H)** Two 3D sections (left and right) through T-zone segmentations, black arrows indicate transverse F-actin bundles running perpendicular on dorsal and ventral surfaces of periphery-orientated MTs. Segmentation colouring is as in Fig. 4A. Scale bars: E-G = 50 nm, H = 100 nm.

**Figure 6.**
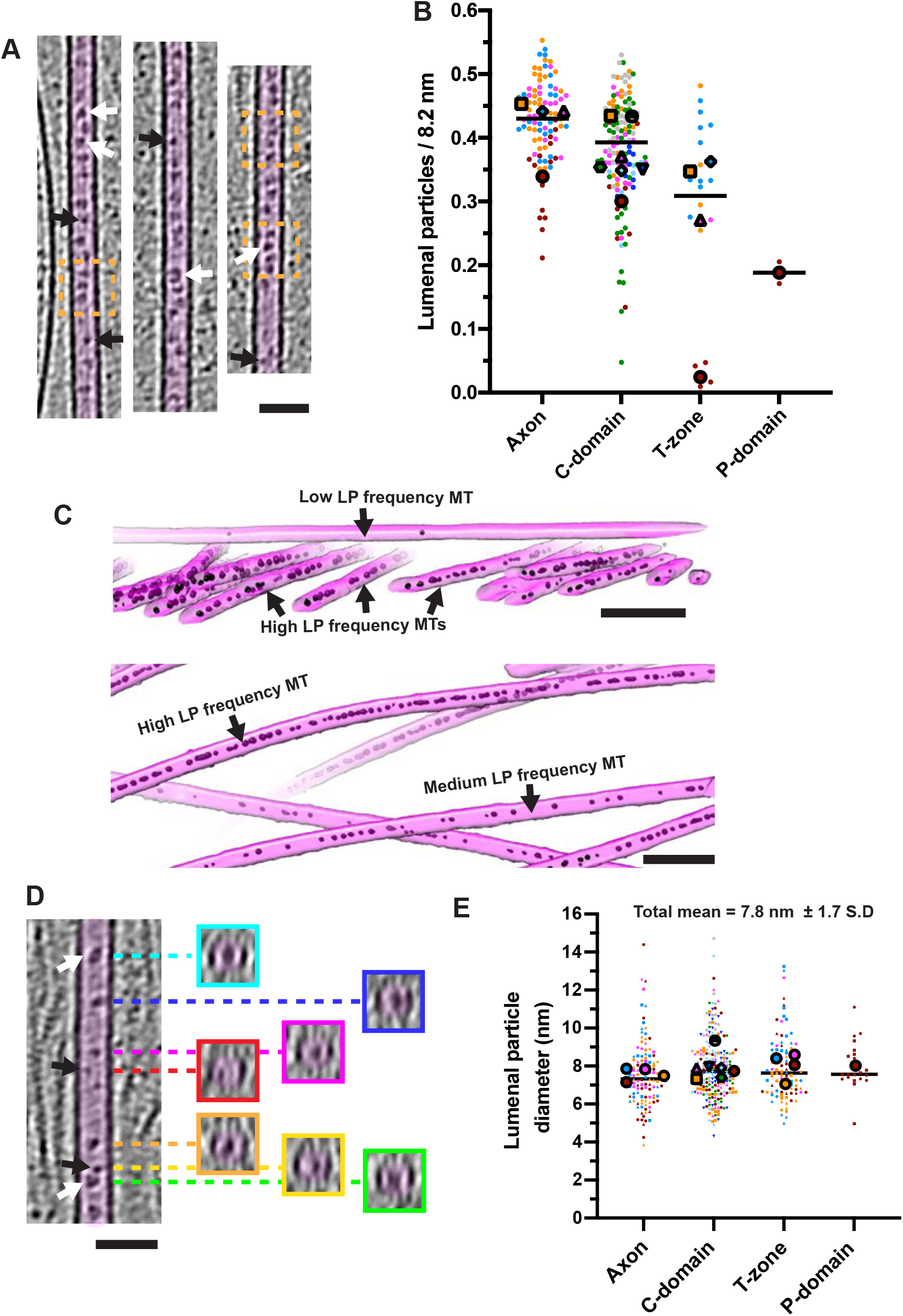
Lumenal particles in neuronal growth cone MTs. **(A)** Longitudinal views (2x binned) of MTs with lumenal particles. Orange dashed boxed regions indicate regions where particles are found every ~8nm. White arrows show larger ring-like particles ~7-10 nm in diameter, black arrows show smaller particles ~3-5 nm in diameter. **(B)** Super-plot of the frequency of lumenal particles per 8.2 nm. Each small data point represents a separate MT coloured by tomogram, mean frequency for individual tomograms are indicated with large coloured shapes, while lines indicate the overall medians. Overall means: Axon = 0.43 ± 0.07 S.D, N = 94 MTs from 4 tomograms; C-domain = 0.38 nm ± 0.09 S.D, N = 134 MTs from 7 tomograms; T-zone = 0.26 ± 0.16 S.D, N = 22 MTs from 4 tomograms; P-domain, = 0.19 ± 0.02 S.D, N=2 MTs from 1 tomogram). **(C)** 3D segmentations of exemplar C-domain MTs (magenta) and lumenal particles (black) illustrating particle frequency variability. **(D)** An MT as in panel A, with transverse sections (~5 nm depth) at positions indicated by the dashed coloured lines. White arrows, larger ring-like particles ~7-10 nm in diameter, black arrows, smaller particles ~3-5 nm in diameter. **(E)** Super-plot of lumenal particle diameters. Each small data point represents a separate lumenal particle coloured by tomogram. Different small data point shapes indicate different MTs within each tomogram. Large coloured shapes indicate mean particle diameters within individual tomograms, lines indicate overall medians. Overall means: Axon = 7.6 nm ± 1.7 S.D, N = 122 from 4 tomograms; C-domain =7.9 nm ± 1.7 S.D, N = 217 from 7 tomograms; T-zone = 7.9 ± 1.6 S.D, N = 97 from 4 tomograms; P-domain = 8.0 ± 1.3 S.D, N = 21 from 1 tomogram. Scale bars: A, D = 50nm; C = 100 nm.

The highest frequency of MT ends as a proportion of MT length was in the T-zone (Fig. 5C) and were mainly blunt-ended, although MTs with tapering ends were occasionally observed (Fig. 5D-F). Of the 6 tapered MT end tomograms, 3 could be determined to be plus ends (axon (2), C-domain (1)) and 1 was a minus end (C-domain) (Fig. 5D, F). No examples of capped MTs were observed, consistent with the absence of γ-tubulin ring complexes from growth cones (Stiess et al., 2010).

In both the T-zone and the P-domain, densities were observed linking the sides of MTs to parallel large axial F-actin bundles, both along their lattice and at their ends (Fig. 5G). In addition, smaller F-actin bundles orientated roughly perpendicular to these MTs were associated with their dorsal and ventral surfaces (Fig. 5H). Our cryo-ET data thereby provide evidence of a diversity of interactions that could mediate integration between the MT and actin filament systems.

### Lumenal particles and lattice defects in growth cone microtubules

The MTs in our tomograms frequently exhibited particles within their lumens, as has been previously described (Atherton et al., 2018; Foster et al., 2021; Garvalov et al., 2006). We refer to these here as “lumenal particles”, rather than MT inner proteins (MIPs,) since in many cases we cannot visualise a direct association with the MT inner wall, a defining characteristic of MIPs (Nicastro et al., 2006) (Fig. 6A). These lumenal particles are not uniformly distributed in the growth cone (Fig. 6B); MTs in the axon and C-domain – where MTs are denser – generally exhibit more lumenal particles (Fig. 6B), whereas they become increasingly sparse in the T-zone and P-domain. However, in some growth cone regions - particularly T-zones - particle density in different tomograms varies substantially (Fig. 6B). Furthermore, within the same tomogram, neighbouring MTs - and even parts of the same MT (Fig. 6A) - exhibit different particle densities (Fig. 6C), with the most densely packed regions containing a particle roughly every 8nm. Particle frequency did not obviously differ between curved and straight MT regions or with proximity to MT ends (Fig. 6C, 5E, Sup. Fig. 2).

The particles themselves are not identical (Fig. 6A,D,E). While the mean lumenal particle diameter is 7.8 nm ± 1.7 (S.D.), the particles range from 3.9 to 14.7 nm, include ring-shaped structures with a diameter of 7-10 nm (Fig. 6D, white arrows) also described by (Foster et al., 2021), as well as smaller, more globular particles of 4-6 nm diameter (Fig. 6D, black arrows). The average size corresponds roughly to a molecular weight of ~ 200 kDa (with a range of 15 to ~1400 kDa, (Erickson, 2009). When MTs are viewed in cross-section, particles are not specifically localised within the lumenal space – some are located centrally while others appear to associate with the inner surface of the MT lattice (Fig. 6D).

We also observed numerous defects and discontinuities in MT lattices in axons and growth cones (Fig. 7A,B). Some were very small, corresponding to one or two (16nm) tubulin dimers, and apparently arising from subunits bending away from the MT axis (Fig. 7B); others were larger (Fig. 7A), with the largest being ~600 nm long. Larger defects were often associated with curved MT regions (Fig. 7A), and thus were more prevalent in the growth cone than the axon (Fig. 7C). No changes in protofilament number, MT architecture or polarity were observed on either side of defects (Fig. 7D), as can occur *in vitro* (Atherton et al., 2018; Chretien and Fuller, 2000; Chretien et al., 1992; Schaedel et al., 2019). Relative to MT length, tomograms of T-zones exhibited the most variable MT defect frequency, with two T-zone tomograms exhibiting an MT defect at least every 2.5 μm of MT length (Fig. 7E). The axon and C-domain exhibited the narrowest range of defect frequency, with a defect being observed every 20 μm on average (Fig. 7E). There was no obvious relationship between the frequency and/or size of defects and the density of the adjacent lumenal particles (Fig. 7A,B).

**Figure 7.**
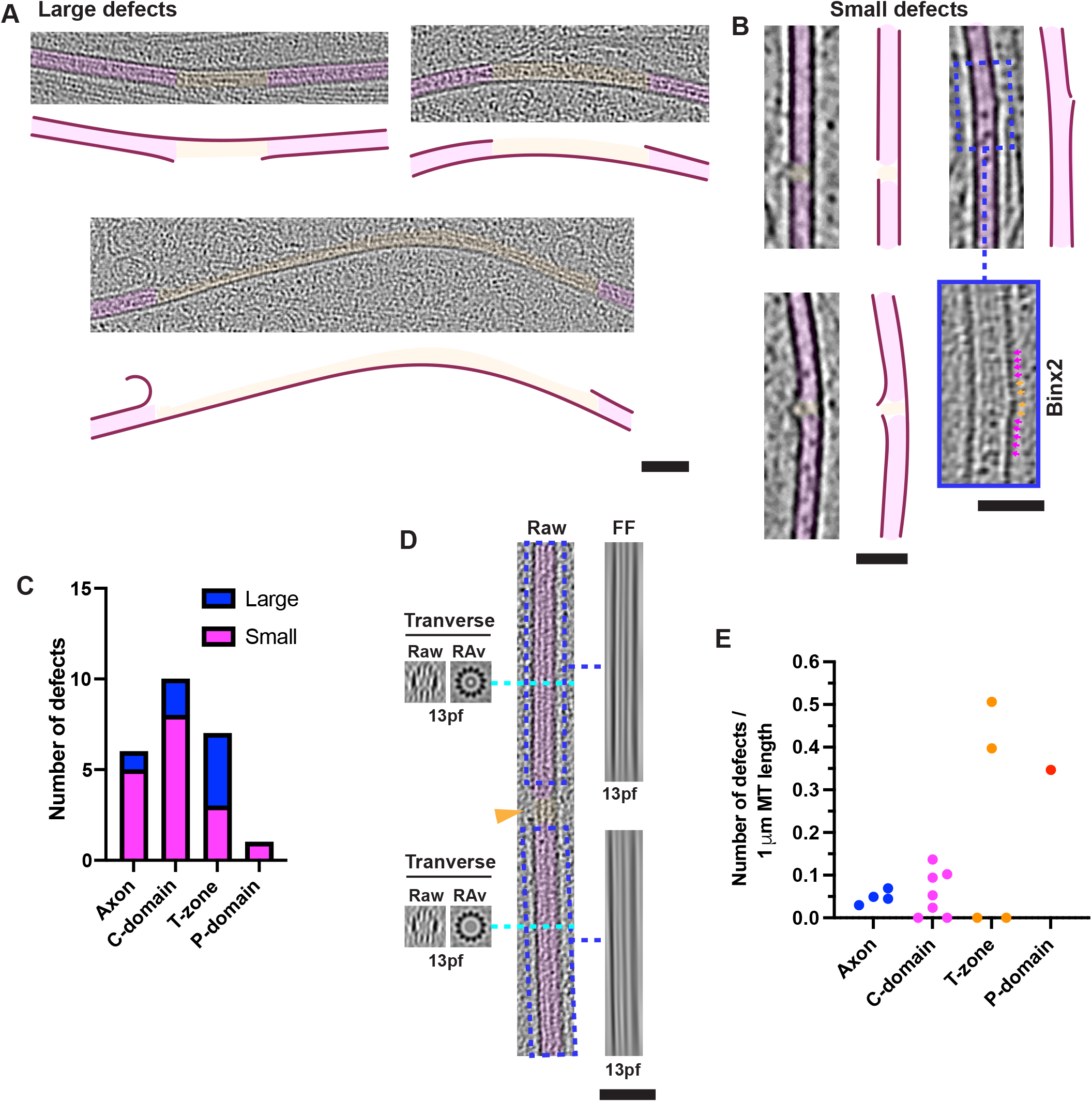
MT lattice defects and discontinuities. **(A)** Longitudinal views of large (> 16 nm long; 30 nm slice, 2x binned) and **(B)** small (< 16 nm long; 10 nm slice, 4x binned) MT lattice defects in growth cones, false coloured in magenta and orange respectively. A traced representation of each MT is shown alongside for clarity. Blue boxed insert in panel B is a 2x binned (5 nm thick) version of the blue dashed boxed region above, with magenta and orange arrows pointing to tubulin monomers in the MT lattice and defect respectively. **(C)** Quantification of the absolute number of small (< 16 nm) and large (> 16 nm) defects (number of tomograms of axon = 4, C-domain = 7, T-zone = 4, P-domain = 1). **(D)** Centre, raw ~30 nm thick longitudinal MT slice including an MT defect (orange arrowhead). Right, Fourier filtered (at the origin, FF) images of the blue dashed boxed regions in the centre panel showing 13-protofilament moiré patterns either side of defect. Left, raw image and rotational average (RAv) of ~10 nm thick MT transverse section, indicated with a cyan dashed line, indicating 13-protofilaments in both cases. **(E)** Frequency of defects per 1 μm MT length. Each data point represents a separate tomogram; axon = 4, C-domain = 7, T-zone = 4, P-domain = 1. Scale bars: A, B, D=50 nm.

### Cryo-ET of Dcx knockout neurons

DCX/Dcx is a modulator of neuronal growth cone morphology and the underlying MT and F-actin organisation (Bielas et al., 2007; Bott et al., 2020; Fu et al., 2013; Jean et al., 2012), and nucleates and stabilises 13-protofilament MTs *in vitro* (Moores et al., 2004). We undertook cryo-ET of hippocampal neuron growth cones from Dcx knockout (KO) mice and analysed their cytoskeletal ultrastructure (Fig. 8, Sup. Fig. 6,7). Like WT neurons, Dcx KO neurons exhibit classical morphology after culture for 2-3 days *in vitro* (Sup. Fig. 6A). By cryo-ET, the general organisation of the pre-cone axon and growth cone in the Dcx KO neurons are indistinguishable from those in WT neurons (Sup. Fig. 6B-D). F-actin ultrastructure and organisation resembles that in WT cells (Sup. Fig. 7A,B) and filopodial F-actin bundles appear indistinguishable from those in WT cells (Sup. Fig. 7B).

**Figure 8.**
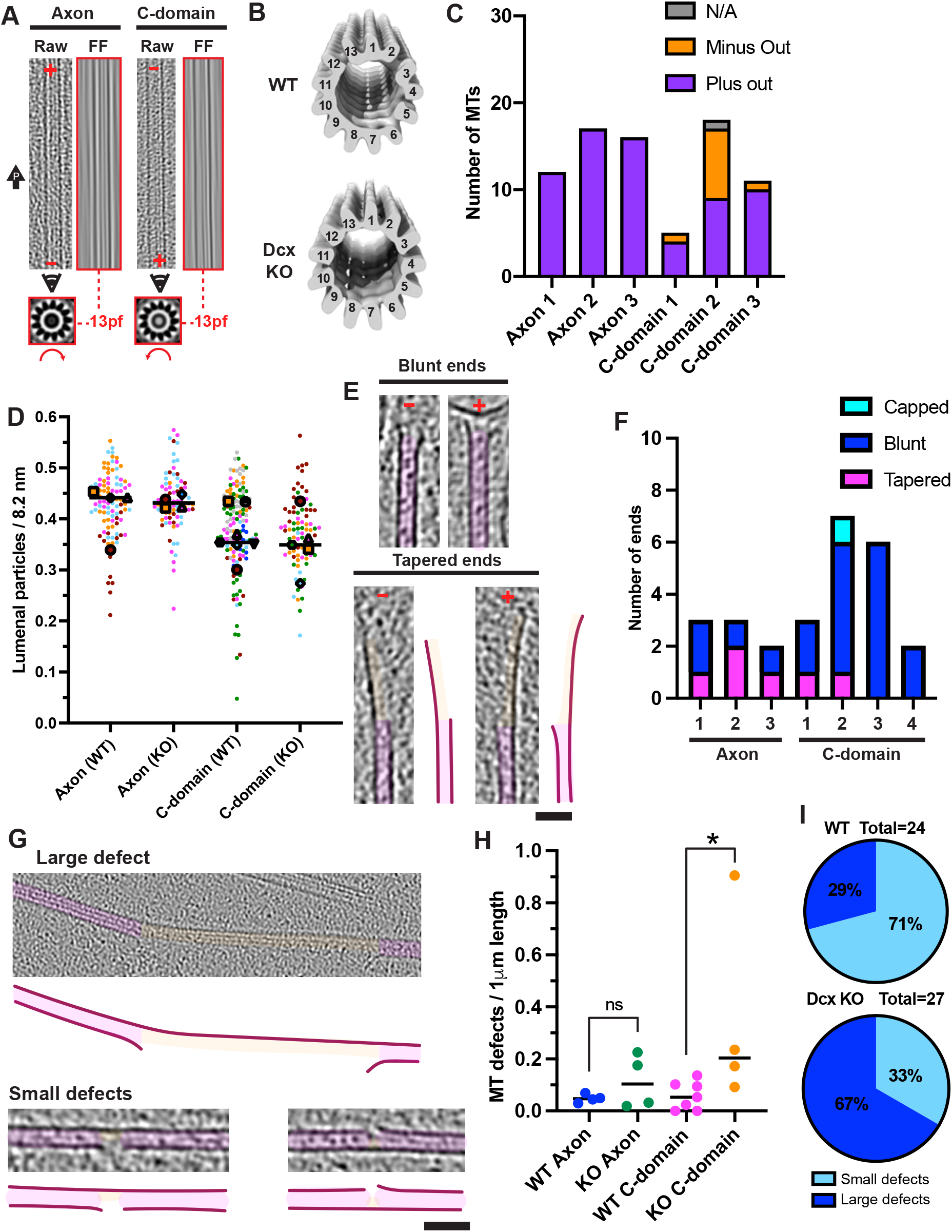
Increased MT defects in doublecortin knock-out neuronal growth cones. ( A) Examplar MTs from Dcx KO neurons showing protofilament number and polarity, as described for Fig 7A. **(B)** Transverse views (tilted, viewed from plus end) of sub-tomogram averages of a random subset of MTs in WT (left, 1200 MT segments from 13 MTs) and Dcx KO (right, 1190 MT segments from 20 MTs) neurons. MT subtomogram averages are built from 13 protofilaments. Volumes were low-pass filtered to their estimated resolutions (Sup. Fig. 7C). **(C)** MT polarity relative to the neuron periphery in individual tomograms. MTs assigned ‘n/a’ were either perpendicular to the axon axis or bent ~180° such that both minus and plus directions were orientated peripherally. Axon = 45 plus-end peripheral MTs, 0 minus-end from 3 tomograms (3 axons), C-domain = 23 plus-end peripheral MTs, 10 minus-end, 1 n/a from 3 tomograms (3 neurons). **(D)** Super-plot of the frequency of lumenal particles per 8.2 nm. Each small data point represents a separate MT coloured by tomogram, mean frequency for individual tomograms are indicated with large coloured shapes, while lines indicate the overall medians. Overall means: WT axon = 0.43 ± 0.07 S.D, N = 94 MTs, 4 tomograms; C-domain = 0.38 nm ± 0.09 S.D, N = 134 MTs, 7 tomograms. Dcx KO axon = 0.43 ± 0.06 S.D, N = 78 MTs, 4 tomograms; Dcx KO C-domain = 0.37 nm ± 0.07 S.D, N = 84 MTs, 5 tomograms. **(E)** Longitudinal views (4x binned) of blunt and tapered MT plus and minus ends in Dcx KO growth cones. **(F)** Quantification of absolute numbers of blunt, tapered and capped MT ends in Dcx KO grow cones. Dataset size is the total number of individual MTs in multiple tomograms (5 Dcx KO axon tomograms, of which 2 had no ends, and 4 Dcx KO C-domain tomograms). **(G)** Longitudinal views (4x binned) of example MT defects in Dcx KO growth cones. **(H)** Frequency of defects per 1 μm MT length in WT and Dcx KO neurons. Each data point represents a separate tomogram; WT axon = 4, WT C-domain = 7, Dcx KO axon = 3, Dcx KO C-domain = 4. Mann-Whitney tests; WT C-domain vs Dcx KO C-domain, p < 0.05 (*), WT axon vs Dcx KO axon, p = 0.89 (not significant). **(I)** Quantification of large and small MT lattice defects as a percentage of total MT lattice defects in WT (left) and Dcx KO (right) pre-cone axons and C-domains. Dataset sizes indicate the total number of defects. Panels E & G: MTs and lattice defects are false coloured in magenta and orange respectively. A traced representation of each MT is shown to its right for clarity. Scale bars: E and G = 50 nm.

MTs in Dcx KO growth cones exhibit exclusively 13-protofilaments, as in WT neurons (Fig. 5A, 8A, Dcx KO, N=79, 7 tomograms); sub-tomogram averages of both WT and Dcx MT segments are also consistent with a 13-protofilament architecture (Fig. 8B). In the axons of Dcx KO neurons, MTs are exclusively oriented with their plus ends towards the growth cone as seen in WT neurons (Fig. 8C, 5B). However, a higher overall proportion of Dcx KO C-domain MTs are oriented with their minus ends towards the neuronal periphery than WT C-domain MTs (29% vs 10%), although this distribution was variable between tomograms (Fig. 8C). Lumenal particle frequency is similar in axons and the C-domain of WT and Dcx KO neurons (Fig. 8D).

MT ends are almost exclusively uncapped (with a possible single exception, Sup. Fig. 7D) in Dcx KO neurons. These MT ends are mainly blunt in appearance, although there appears to be a moderate increase in the proportion of tapered MT ends in the Dcx KO neurons (Fig. 7E,F vs. Fig. 5D). Importantly, however, a higher frequency of MT defects were observed in Dcx KO C-domains (Mann-Whitney test, p<0.05) compared with WT C-domains (Fig. 8G-H) with, in one C-domain tomogram a defect observed for every 1 μm of polymer length (Fig. 8H). Furthermore, a higher proportion of defects found in Dcx KO growth cones are > 16nm (Fig. 8I).

## DISCUSSION

Cryo-ET, neural network-driven density segmentation, quantitative analysis and sub-tomogram averaging revealed the distinct specialisations of the cytoskeleton in different parts of neuronal growth cones, reflecting the complexity of their behaviour and regulation. Within the tightly bundled actin filaments that dominate the P-domain (Fig. 3C, D), different types of short F-actin cross-linkers - which our data cannot discriminate - likely coexist. However, bundle architecture is most consistent with the involvement of fascin (Aramaki et al., 2016; Ishikawa et al., 2003; Jansen et al., 2011; Kureishy et al., 2002; Winkelman et al., 2016; Yang et al., 2013), and with fascin’s known localisation and role in filopodia formation (Cohan et al., 2001; Sasaki et al., 1996). *In vitro*, fascin-mediated actin bundles with very similar architecture exhibit a maximum size of ~20 filaments (Claessens et al., 2008), limited by bundle packing. However, we and others (Aramaki et al., 2016) observe examples of much bigger bundles (> 100 filaments) within filopodia. We also observe flexible actin filaments weaving between bundles which, together with other actin modulators, may loosen bundle packing sufficient to allow larger overall bundles. Larger but less stiff bundles may be important for the balance between the dynamic activities of filopodia and shape maintenance.

Alongside the filopodia-derived bundles in the P-domain and T-zone, we also observed shorter-pitch actin filaments. The ~27 nm half helical pitch length, wider appearance and 3D reconstruction of these filaments suggest that cofilin is bound, consistent with its localisation to the P-domain and roles in filopodial F-actin severing and recycling (Bamburg and Bray, 1987; Breitsprecher et al., 2011; Gungabissoon and Bamburg, 2003; Marsick et al., 2010). *In vitro*, cofilin binds cooperatively to F-actin, alters the actin filament structure to a half helical pitch length of ~27 nm, and promotes ADP-actin filament severing (Bobkov et al., 2004; Galkin et al., 2001; Huehn et al., 2020; McCullough et al., 2008; McGough et al., 1997; Tanaka et al., 2018). When associated with bundles in our cryo-ET data, these short-pitch cofilin-like filaments were frequently peripheral, exhibited more positional fluidity within the bundle, and often detached from the main filament cluster. They were also not regularly cross-linked to long-pitch filaments, likely due to mismatch between helical pitches. Our data provide good evidence that well-decorated cofilin-bound actin filaments exist in the P-domain, and suggest that cofilin decoration may enable filament detachment from bundles before severing, thereby contributing to overall retrograde flow from the front of the cell (Vitriol et al., 2013).

Despite the cofilin-type activity in the P-domain, filopodia derived bundles ultimately taper and terminate in the T-zone, often with bundles splitting apart and bending away into actin networks that run roughly perpendicular to filopodia bundles. How filaments are bent away from F-actin bundles is unclear; however myosin-II is known to unbundle F-actin, contribute to severing, and produce retrograde flow and bending forces in the T-zone (Burnette et al., 2008; Ishikawa et al., 2003; Medeiros et al., 2006; Schaefer et al., 2002). Further, since filament peeling is observed in both directions from filopodia-derived bundles, the resulting anti-parallel networks necessary for T-zone myosin-II-mediated contractility can be formed. Consistent with this, myosin-II/F-actin *in vitro* complexes share striking similarities to the tapered unbundling observed here *in situ* (Haviv et al., 2008).

The density of T-zone F-actin networks suggests that they do not simply create a physical barrier to MTs entering the P-domain. However, close association of MTs with F-actin is consistent with previously observed restriction of MT entry into the growth cone periphery primarily by retrograde flow (Burnette et al., 2008; Burnette et al., 2007; Kahn and Baas, 2016; Schaefer et al., 2002; Schaefer et al., 2008). Retrograde flow has also been shown to bend and break growth cone MTs, thereby contributing to their overall dynamics (Waterman-Storer and Salmon, 1997). However, a number of end-binding proteins, MAPs and motors important for neuronal migration form links to F-actin in growth cones (Cammarata et al., 2016; Geraldo and Gordon-Weeks, 2009; Kahn and Baas, 2016), and there is abundant evidence in our data of such MT-F-actin cross-linkers which likely mediate cytoskeleton cross-talk (Fig. 5G).

Growth cone MTs contain diverse and unevenly distributed lumenal particles. As MTs in the periphery are more dynamic compared to the axon and C-domains, MTs with high and low lumenal particle densities may represent populations of stable and dynamic MTs respectively; the more even distribution of lumenal particles in axon MTs is consistent with this idea (Foster et al., 2021). The identities of these lumenal particles remain unclear - some cytoplasmic components may simply be trapped during MT polymerisation. However, other densities are likely to be functionally relevant - for example, the neuronal MAP6 is likely to be among the components forming these particles (Cuveillier et al., 2020). Based on the volume of the MT lumen, the average volume of the particles and their average frequency, we estimate that lumenal particles occupy roughly 7% of the total inner volume of neuronal MTs. However, their precise roles are unclear, and the MT lumenal space remains an intriguing and poorly understood cellular compartment.

At least a subset of growth cone MTs are dynamic (Baas and Black, 1990; Edson et al., 1993; Tanaka et al., 1995). Strikingly, however, the large majority of MT ends we observed are relatively blunt, a few exhibit longer sheet-like extensions, and we found no examples of extensively curled protofilaments that have been proposed to characterise both growing and shrinking MTs (McIntosh et al., 2018). The large number of neuronal MAPs may constrain the morphologies of dynamic MT ends such that curling protofilaments are less likely to form. MT lattice defects, and the repair processes they promote, also contribute to regulation of MT dynamics (Thery and Blanchoin, 2021). In our data, MTs throughout the growth cone exhibit discontinuities or substantial breaks, particularly in the T-zone. Lumenal particleforming MAP6 induces MT defects *in vitro* (Cuveillier et al., 2020); however, in our data there is no clear correlation between defects and lumenal particle density, and defects are more common in peripheral regions with overall fewer particles. Large defects were frequently observed on curved MTs, and it therefore seems likely that at least some larger defects arise from strain-induced MT fraying, while short MTs may represent remnants of broken MTs caused by retrograde flow (Burnette et al., 2008; Schaefer et al., 2002; Schaefer et al., 2008; Waterman-Storer and Salmon, 1997).

DCX/Dcx has a number of reported roles in migrating neurons related to its cytoskeleton regulation functions (Bielas et al., 2007; Deuel et al., 2006; Friocourt et al., 2003; Fu et al., 2013; Jean et al., 2012; Li et al., 2021; Tint et al., 2009). In Dcx KO neurons, growth cone architecture and the overall organisation and distribution of the cytoskeleton were comparable to WT neurons; this included F-actin organisation, in contrast to an earlier report (Fu et al., 2013). *In vitro*, whereas MTs polymerised from diverse tubulin sources including mouse brain tubulin (Vemu et al., 2017) polymerise with a range of protofilament architectures (Ray et al., 1993), DCX nucleates and selectively stabilises 13-protofilament MTs (Moores et al., 2004). In the Dcx KO growth cones, however, all MTs exhibited a 13-protofilament architecture as in the WT neurons, highlighting the role of diverse cellular factors in addition to Dcx in ensuring uniformity of MT architecture even in the absence of γ-TuRCs (Stiess et al., 2010). However, in our Dcx KO tomograms, C-domain MTs exhibited a higher proportion of minus end out MTs, a higher frequency of defects, and a higher proportion of large compared to small defects. DCX/Dcx is highly expressed in growth cones where it has been shown to stabilise and straighten MTs (Jean et al., 2012; Tint et al., 2009) and its binding is modulated by MT curvature (Bechstedt et al., 2014; Ettinger et al., 2016). Taken together with our data, this suggests that absence of Dcx may lead to a higher frequency of curvature-induced MT fraying or breakage in C-domains, where Dcx is particularly abundant (Sup. Fig. 1B), and greater persistence of MT defects overall.

How might this potential role for DCX/Dcx explain brain developmental phenotypes caused by its absence? Immature neurons experience substantial compression forces during navigation through the layers of the developing brain (Tyler, 2012); the requirement to maintain and repair ruptured MTs during migration may be particularly acute, and DCX/Dcx may be involved in this process. Cryo-ET will continue to be a vital tool for the molecular visualisation and dissection of machinery involved in key cellular processes including responses to guidance signals and injury in neurons. Insights arising from such studies will also inform developments of new treatments for neuronal damage and degeneration (Griffin and Bradke, 2020).

## MATERIALS AND METHODS

### Transgenic mice and genotyping

WT and Dcx KO mice were maintained on the Sv129Pas background with more than ten generations of backcrosses (Kappeler et al., 2007; Kappeler et al., 2006). Genotyping was performed by PCR to verify the inactivation of the Dcx gene in KO animals (Kappeler et al., 2006). All experiments were performed in accordance with institutional, national and international guidelines (EC directive 2010/63/UE) and were approved by the local ethical committees (French MESR N°: 00984_02). The day of confirmation of vaginal plug was defined as embryonic day zero (E0.5).

### Neuronal cell dissociation and isolation

Embryonic mouse hippocampus was dissected at E17.5 in ice-cold 0.02 M HEPES in Ca^2+/^Mg^2+^-free HBSS (Gibco) and primary cell suspensions dissociated via incubation with 2.5 mg/ml trypsin (Sigma). After centrifugation to a pellet (5 min, 300g), cells were resuspended in culture media for light or electron microscopy experiments as indicated below.

### Cell culture for fluorescence microscopy

For fluorescence microscopy experiments dissociated neuronal pellets were resuspended and cultured in Neurobasal medium supplemented with B27 (1%, Gibco), GlutaMAX (2 mM, Gibco), penicillin (100 units/ml, Invitrogen) and streptomycin (100 mg/ml, Invitrogen). Only mouse litters comprised of at least 1 WT and 1 Dcx KO animal were used. Dissociated cells were plated on 14 mm-diameter coverslips (0.3 x 10^5^ cells/ coverslip) coated with poly-L-lysine (0.05 mg/ml, Sigma) and laminin (0.01 mg/ml, Invitrogen) and cultured at 37°C in the presence of 5% CO_2_.

### Fluorescence microscopy

Neurons were fixed after 3 days in vitro (DIV) following the protocol from (Tint et al., 2009), applying PEM buffer with 0.3% gluteraldehyde, 0.1% IgepalCA630, and 10 μM taxol for 10 min R.T. Coverslips were then washed 5X with PBS, 1X 15 min in PBS with 0.5% triton, 2X 7 min with 10 mg/mL sodium borohydride in PBS, and 5X with PBS. Neurons on coverslips were incubated with blocking solution (5% normal goat serum, 0.1% Triton X-100 in 1X PBS) for 1 h at room temperature (RT), followed by incubation with primary antibodies for Dcx (Francis et al., 1999) and neuron-specific β3-tubulin (TUJ1, RD Biosystems) in blocking solution for 2 h at R.T. or overnight at 4°C. Secondary antibodies (Invitrogen) were incubated in blocking solution combined with Hoechst for 60 min at RT in the dark, followed by 60 min incubation with phalloidin-A633 (Invitrogen) in PBS. Coverslips were mounted with Fluoromount G (Southern Biotechnology).

### Neuronal cell culture and preparation for cryo-EM

For electron microscopy experiments dissociated neuronal pellets were resuspended in Hibernate-E medium (Gibco) supplemented with B27 (Invitrogen) and Pen-Strep (100 units/ml penicillin and 100 mg/ml streptomycin, Invitrogen) and shipped overnight from Paris to London in the dark at 4°C in Hibernate-E media. Quantifoil R2/4 gold G200F1 Finder EM grids were placed on the surface of plastic culture dishes and sterilized under UV light. The dishes and overlaid EM grids were then coated with poly-d-lysine (Millipore) and mouse laminin (Thermofisher). Within 24–30 h of initial dissociation neurons were pelleted by centrifugation (5 min, 300g), then resuspended and plated in Neurobasal medium with B27 and Pen-Strep and cultured at 37°C in the presence of 5% CO_2_ for 2–3 d *in vitro* without media change until the characteristic morphology was observed using phase contrast light microscopy. Grids were extracted from culture dishes, 3 μl of conditioned culture media added, blotted for 8-10s and plunge-vitrified in liquid ethane using a Vitrobot (FEI) set at 37°C and 80% humidity.

### Cryo-ET data collection

Cryo-electron tomography was performed using a Tecnai G2 Polara operating at 300kV with a Quantum post-column energy filter (Gatan), in zero-loss imaging mode with a 20 eV energy-selecting slit. Single-axis cryo-electron tomography was performed at 3-6 μm defocus with a K2 Summit direct electron detector (Gatan) operating in counting mode (at 5 e-per pixel per second) with a final sampling of 5.38 Å per pixel. Dose-symmetric tilt-series (Hagen et al., 2017) within a −60° to +60° tilt range were collected at 3° increments to give a total dose of 110 e-/Å^2^. Movies of three or four subframes per second were collected at each tilt and aligned on-the-fly using MotionCor2 (Zheng et al., 2017). 30 or 16 tilt series of a ~2.15 by 2.15 μm field of view were collected from different WT (derived from 12 neurons, 7 different mice and 10 different grids) and Dcx KO (derived from 9 neurons, 4 different mice and 6 different grids) growth cone regions, that lay over 2 μm diameter holes in the carbon substrate of the EM grid (Sup. Fig. 1C).

### Cryo-ET image processing

Fiducial-less alignment via patch-tracking was performed on each tilt series in the Etomo graphical user interface to IMOD v.4.9.0 (Kremer et al., 1996). CTF was determined at each tilt using CTFFIND4 (Rohou and Grigorieff, 2015) and then a dose-weighted aligned tilt-series was produced using Summovie (Grant and Grigorieff, 2015) under the control of custom scripts. Three-dimensional CTF correction and tomogram reconstruction was performed by weighted back-projection of dose-weighted tilt series with novaCTF (Turonova et al., 2017) using custom scripts.

Semi-automated tomogram segmentation was performed using the tomoseg module of EMAN2 v.2.2 (Chen et al., 2017). 4x binned tomograms with low frequencies amplified via a positive B-factor of 100,000 were used for training and segmentation purposes. Segmentations were further cleaned manually in Chimera (Pettersen et al., 2004) for display, via a combination of masking, ‘volume eraser’ and ‘hide dust’ tools.

Subtomogram averaging of MT and F-actin filaments was performed in Dynamo. 341 volumes of 68 by 68 by 68 nm were picked from 4x binned tomograms using filament model tools. Initial 40 Å low-pass filtered references were generated from data aligned only along filament axes. Initial coarse alignment was performed using 4x binned data, before finer alignment with 1x binned data.

## ACKNOWLEDGEMENTS

M.A.S. and F.F. are associated with the BioPsy Labex project and the Ecole des Neurosciences de Paris Ile-de-France (ENP) network and thank the IFM animal experimentation platform for mouse maintenance and production (supported by Inserm, the Région Ile de France and the Fondation Bettencourt Schueller). We thank Natasha Lukoyanova for support during cryo-EM data collection, and Joshua Hutchings, Joseph Beton and Phillip Gordon-Weeks for useful discussions and technical advice.

