## Supporting Information for "Visualising the cytoskeletal machinery in neuronal growth cones using cryo-electron tomography"

**Supplementary Figure 1. Cryo-electron tomography of primary neuronal growth cones and axons.**

**(A)** Mouse primary hippocampal neuron cultured for 3 days *in vitro*, demonstrating the characteristic polarised stage 3 morphology, labelled as indicated with DAPI (nucleus) and phalloidin (F-actin) dyes and antibodies to β3-tubulin and Dcx; **(B)** Fluorescence light microscopy images of a mouse hippocampal neuron growth cone labelled as indicated with phalloidin (F-actin) dye and antibodies to β3-tubulin and Dcx; the central domain (C), transition zone (T) and peripheral domain (P) are indicated on the composite image. **(C)** Low magnification cryo-EM image of a growth cone. The cell membrane is depicted in blue; the C-domain (C), T-zone (T) and P-domain (P) are indicated; F, filopodia; L, Lamellipodia; H, underlying holes in the carbon substrate; I, Ice contamination. **(D)** Example tomogram and corresponding segmentation from an axonal region. Left panel shows central section through a binned x 4 tomogram, with MTs (magenta), large vesicular organelles (green) and outer membrane (blue) indicated with overlaid false colouring. Direction of the leading edge is indicated with large black arrows containing a 'P' and edges of the carbon substrate hole are indicated with small black arrows. The middle panel shows the corresponding whole volume 3D semi-automated segmentations. The right panel shows a transverse section of the segmented region indicated with a dashed line in the corresponding central panel (section viewing direction also indicated in central panel). Segmentation false colouring is according to the key. Scale bars: (A) = 20 μm, (B) = 10 μm, (C) = 2 μm, D = 200 nm.

**Supplementary Figure 2. The molecular landscape of neuronal growth cones.**

Example tomograms and corresponding segmentations from different growth cone regions; **(A)** C-domain, **(B)** T-zone, **(C)** P-domain, **(D)** P-domain filopodia. Left panels show central sections through binned x 4 tomograms, with MTs (magenta) and outer membrane (blue) indicated with overlaid false colouring. Direction of the leading-edge distal periphery is indicated with large black arrows containing a 'P' and edges of the carbon substrate hole are indicated with small black arrows. Middle panels show corresponding whole volume 3D semi-automated segmentations. Right panels show transverse sections of the segmented regions indicated with dashed lines in corresponding central panels (section viewing direction also indicated in central panels). Segmentation colouring is according to the key. Scale bars = 200 nm.

**Supplementary Figure 3. Ultrastructure of the ordered P-domain F-actin arrays.** **(A)** Super-plot (Lord et al., 2020) of maximum filament number in width and height of P-domain bundles. Each data point represents an individual filopodial F-actin bundle (N = 14 from 4 tomograms, shown as different colours), with a line indicating the overall median. Mean values for each tomogram are shown in their respective colours with larger shapes. Overall mean width, 11.3 ± 2.2 S.D, mean height 6.1 ± 1.4 S.D filaments. **(B)** Super-plot of inter-filament distances in P-domain F-actin bundles. Each data point represents a separate adjacent filament pair (N = 23 from 3 tomograms, shown as different colours). Mean values for each tomogram are shown in their respective colours with larger shapes. Line indicates the overall mean (12.4 nm ± 1.3 S.D). **(C)** Super-plot of regular longitudinal crosslink separation distance in P-domain F-actin bundles. Each data point represents an individual pair of adjacent cross-links along the F-actin longitudinal axis (N = 25 from 3 tomograms, shown as different colours). Mean values for each tomogram are shown in their respective colours with larger shapes. Line indicates the overall mean (37.9 nm ± 2.1 S.D).

**Supplementary Figure 4. Resolution estimates for sub-tomogram averages of growth cone F-actin structures**

Fourier Shell Correlation (FSC) curves generated from half sets for subtomogram-averages of short-pitch F-actin filaments (grey), long-pitch F-actin filaments (green) and hexagonal arrays of P-domain F-actin bundles (yellow). Resolutions are indicated at the FSC = 0.5 criterion.

**Supplementary Figure 5. Characteristics of short-pitch F-actin in the P-domain and T-zone**

**(A)** i) Longitudinal and ii) transverse (~20 nm depth) views of a large F-actin bundle in a 2x binned P-domain tomogram. Dashed cyan line in i) illustrates position of transverse section in ii). Examples of ~37 nm F-actin half helical pitch lengths are indicated with yellow dashed arrows, while shorter, rarer ~27 nm F-actin half helical pitch lengths are shown with green dashed arrows. A short-pitch filament is false coloured in green and additionally indicated in ii) with a green arrowhead. In this and subsequent longitudinal views, more peripheral and more central regions of the neuron are at the top and bottom of the images respectively. **(B)** Longitudinal view of a large F-actin bundle in a 2x binned P-domain tomogram, with two short-pitch filaments false coloured in green. F-actin half helical pitch lengths are indicated with arrows as in panel A. **(C)** Longitudinal view of a large F-actin bundle in a 2x binned P-domain tomogram, showing a single filament transitioning from short to long-pitch (with short and long-pitch filament segments false coloured in green and yellow respectively). F-actin half helical pitch lengths are indicated with arrows as in panel A. **(D)** Longitudinal view of an individual non-bundled short-pitch F-actin filament in a 2x binned T-zone tomogram, false coloured in green. F-actin half helical pitch lengths are indicated with arrows as in panel A. Scale bars: A - D = 40 nm.

**Supplementary Figure 6. Cryo-electron tomography of primary neuronal growth cones and axons from Dcx KO mice.**
**(A)** Mouse Dcx KO primary hippocampal neuron cultured for 3 days *in vitro*, demonstrating the characteristic polarised stage 3 morphology, labelled as indicated with DAPI (nucleus) and phalloidin (F-actin) dyes and β3-tubulin antibody. **(B)** Low magnification example cryo-EM image of a Dcx KO growth cone. The cell membrane is depicted in blue; the C-domain (C), T-zone (T) and P-domain (P) are indicated; F, filopodia; L, Lamellipodia; H, underlying holes in the carbon substrate; I, Ice contamination; a representative region for tilt series data collection is indicated by the dotted turquoise boxed region. **(C, D)** Example tomogram and corresponding segmentation from a Dcx KO axon **(C),** C-domain **(D)**. Left panel shows central section through a binned x 4 tomogram, with MTs (magenta), large vesicular organelles (green) and outer membrane (blue) indicated with overlaid false colouring. Direction of the leading edge is indicated with large black arrows containing a 'P' and edges of the carbon substrate hole are indicated with small black arrows. The middle panels show the corresponding whole volume 3D semi-automated segmentations. The right panels show a transverse section of the segmented region indicated with a dashed line in the corresponding central panel (section viewing direction also indicated in central panel). Segmentation false colouring is according to the key. Scale bars: A = 20 μm, B = 2 μm, C, D = 200 nm.

**Supplementary Figure 7. Normal ultrastructural organisation of filopodial F-actin networks in Dcx KO growth cones.**

**(A)** Example tomogram and corresponding segmentation from filopodia in a Dcx KO P-domain. Left panel shows central section through a binned x 4 tomogram, with outer membrane (blue) indicated with overlaid false colouring. Direction of the leading edge is indicated with large black arrows containing a ‘P’. The middle panel shows the corresponding whole volume 3D semi-automated segmentation. The right panel shows a transverse section of the segmented region indicated with a dashed line in the corresponding central panel (section viewing direction also indicated in central panel). Segmentation false colouring is according to the key. **(B)** Transverse views through filopodial F-actin bundles (~20 nm depth) in WT and Dcx KO P-domains, showing their similar characteristic hexagonal arrangement. **(C)** Fourier Shell Correlation (FSC) curves generated from half sets of subtomogram-averages of MTs from WT (blue) and Dcx KO (green) neurons. Resolutions are indicated at the FSC = 0.5 criterion. **(D)** Longitudinal view of a potential capped MT minus end in a Dcx KO neuron, shown as a 10 nm thick slice through a 4x binned tomogram. The MT is false coloured in semi-transparent magenta. The minus end is indicated with a red ‘–’. Scale bars: A = 200 nm, B = 10 nm, C = 50 nm.

### Supplementary Figure 1

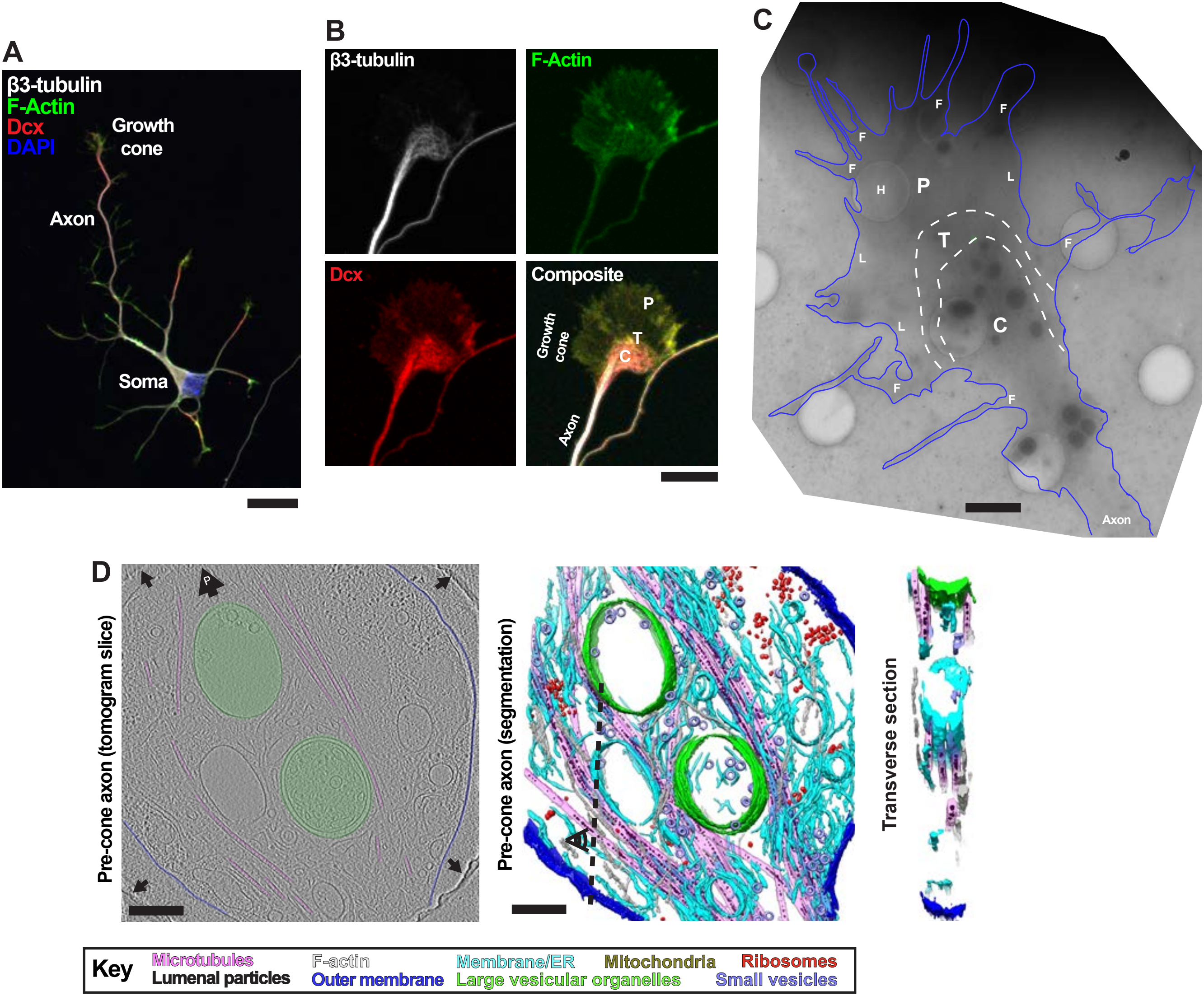

### Supplementary Figure 2

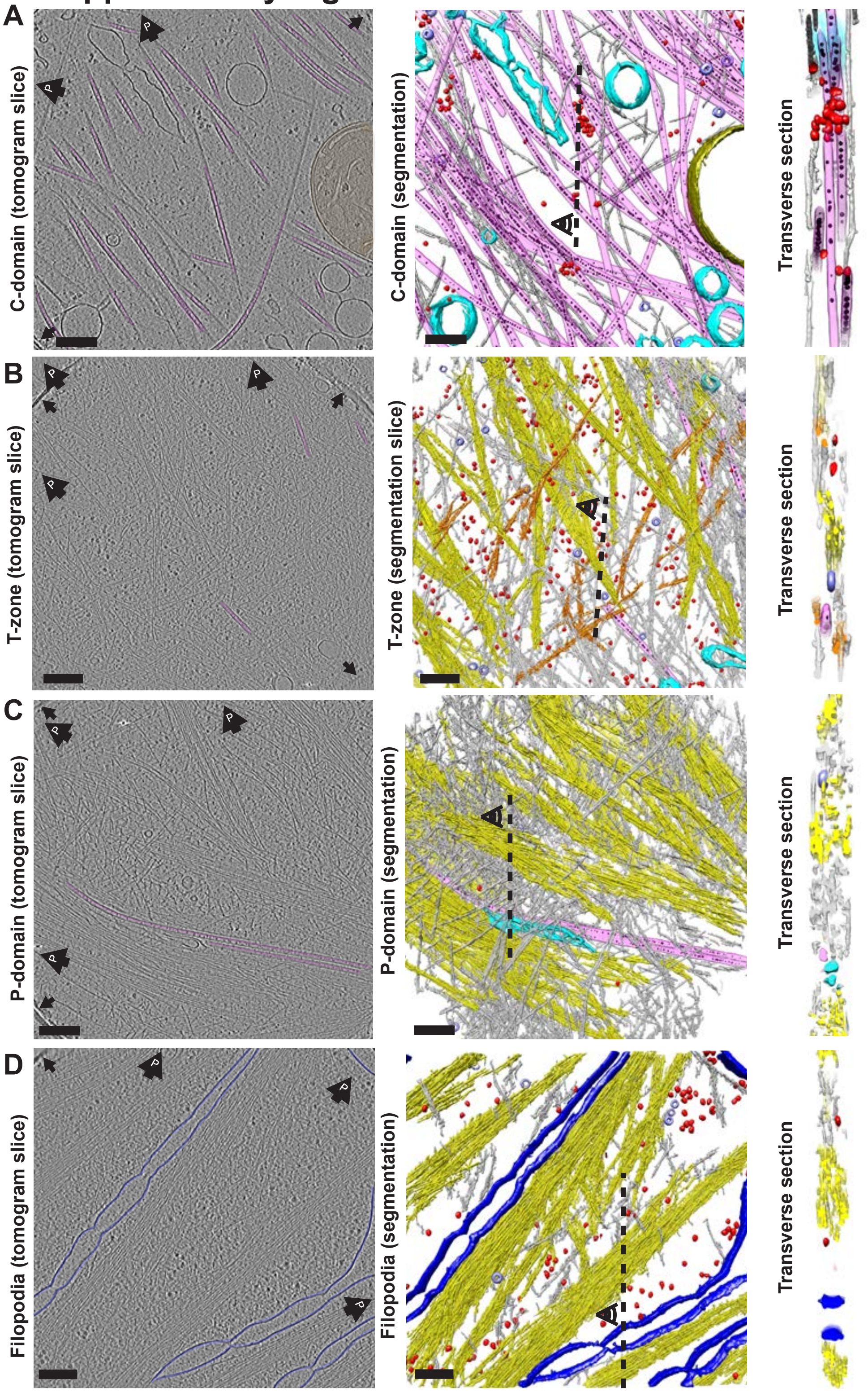

### Supplementary Figure 3

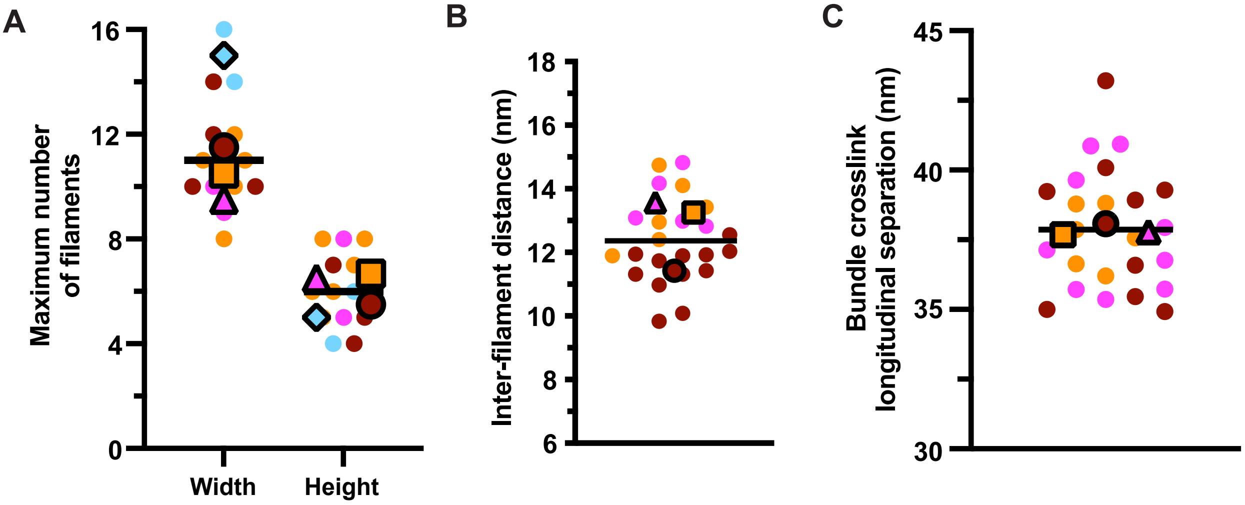

### Supplementary Figure 4

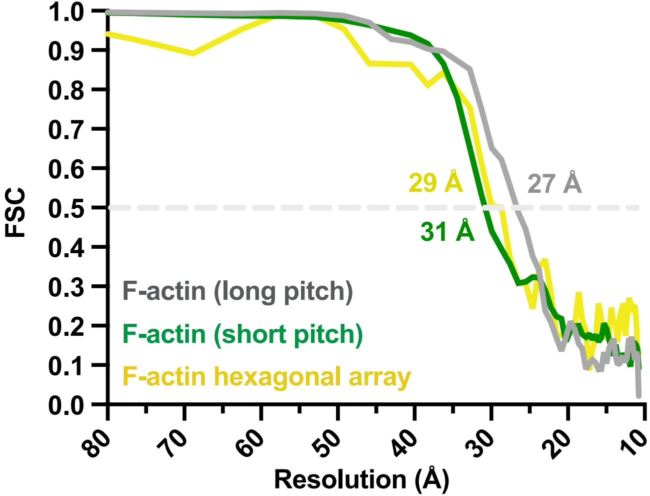

### Supplementary Figure 5

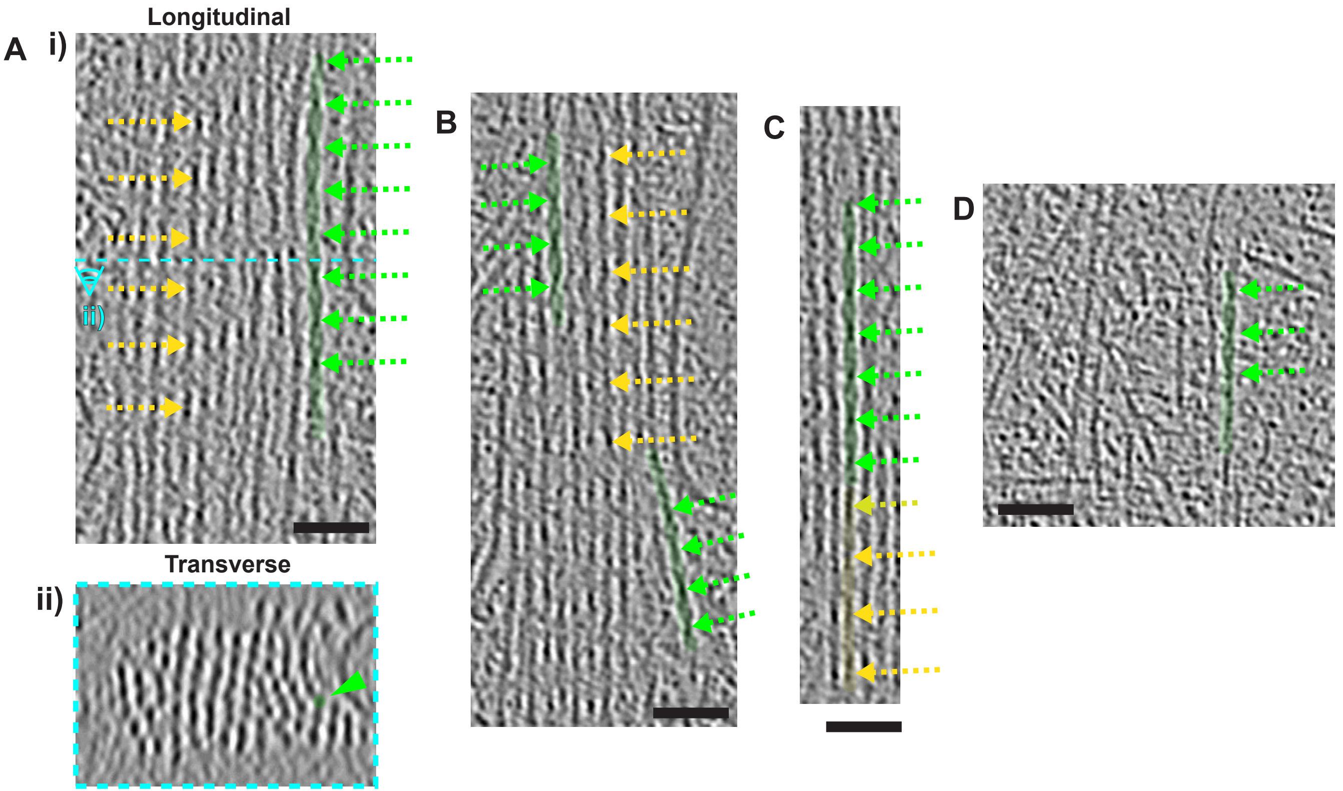

### Supplementary Figure 6

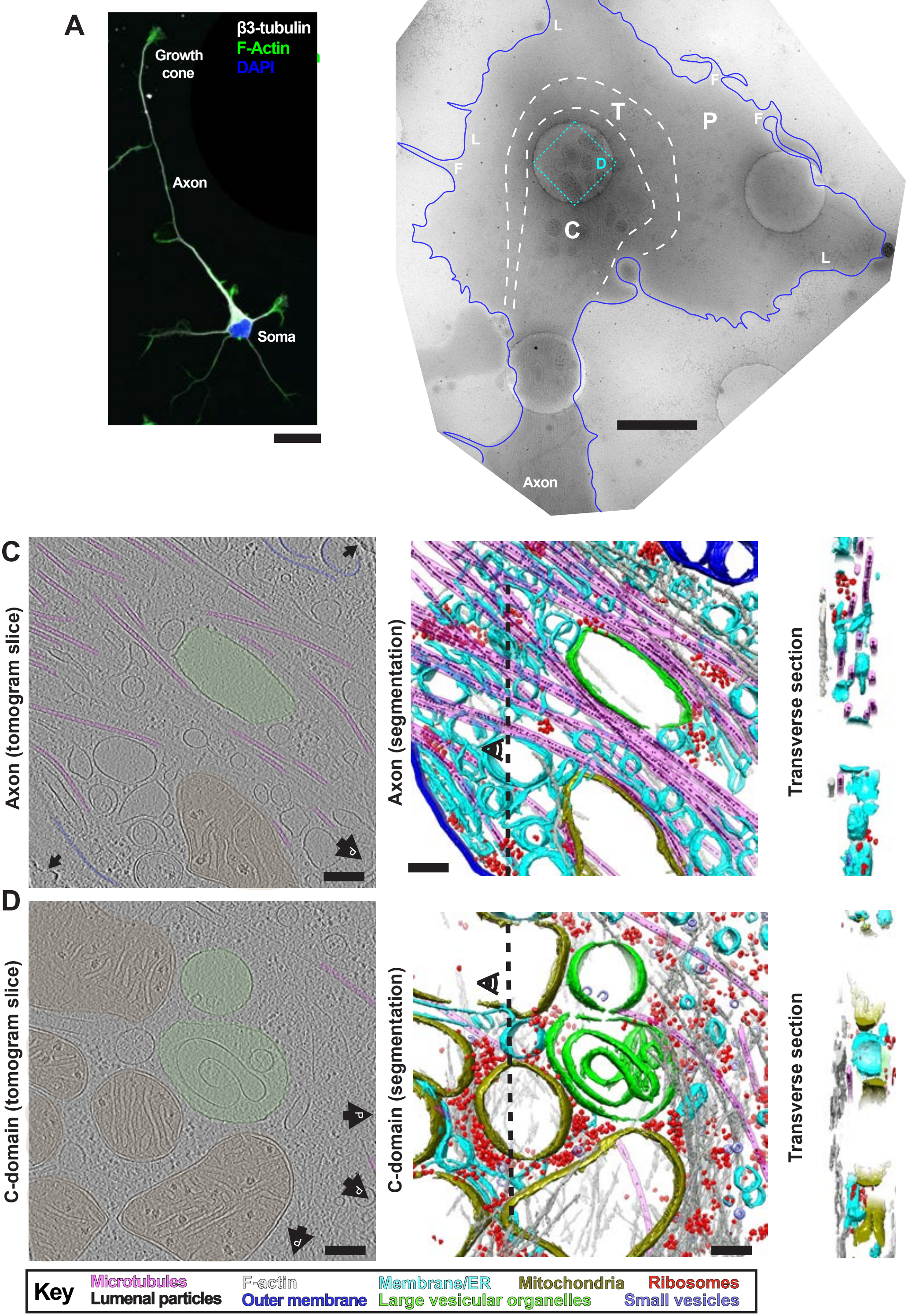

### Supplementary Figure 7

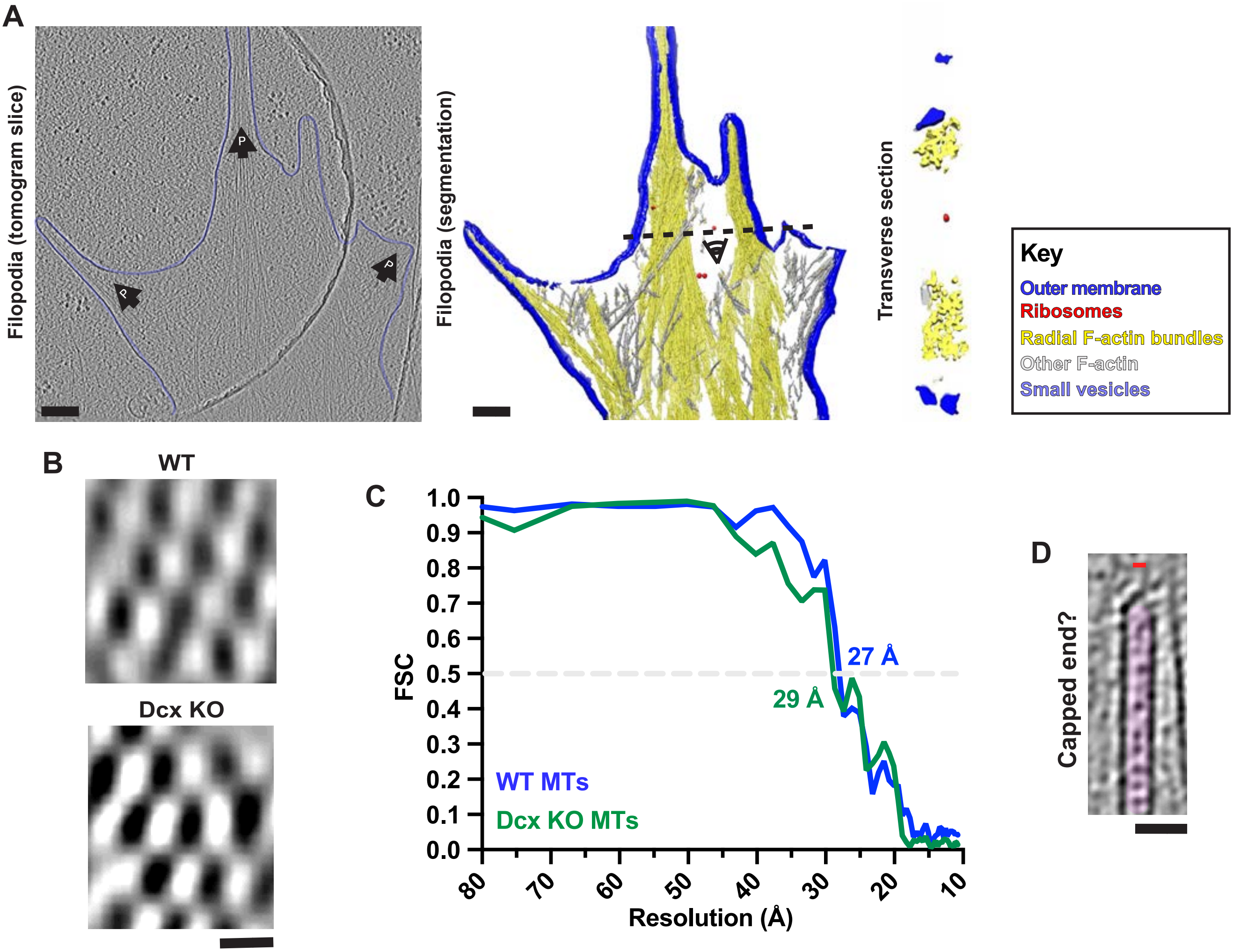

**Supplementary Video 1. Overview of neuronal growth cone cryo-electron tomography data and segmentation.**
Overview of cryo-ET procedures used during this study. The video starts with a low-magnification electron micrograph of a neuronal growth cone, with the cell membrane depicted in blue; the C-domain (C), transition zone (T) and peripheral domain (P) and ice contamination (I) indicated. A representative region over a hole in the carbon for tilt series data collection is indicated by the dashed cyan box. The video then shows the corresponding tilt series acquired for this region, with the peripheral leading edge of the growth cone indicated. Next, the corresponding reconstructed 2x binned tomogram is shown, followed by the corresponding semi-automated segmentation with features coloured as indicated.

**Supplementary Video 2. Fluidity within radial F-actin bundles.**
Views though a radial F-actin bundle (as indicated in yellow false colour) in transverse sections (~20 nm depth), showing filaments in a rough hexagonal stacking, yet with a degree of fluidity with individual filaments being lost, added or moving position within the bundle. Scale bar = 25 nm.

**Supplementary Video 3. Sub-tomogram average of hexagonal arrays in radial F-actin bundles.**
The 3D sub-tomogram average (yellow mesh) calculated from hexagonal arrays of P-domain F-actin bundles. The video starts by showing a transverse section of the masked hexagonal bundle, with the central and surrounding filaments numbered in yellow (0 and 1-6 respectively). Cross-links from the central filament, indicated as coloured balls to help visualisation (colouring as indicated), are also numbered in grey according to the filaments connected by each cross-link. The video then clips through in the transverse direction, and rotates the volume from a longitudinal view, which is then rotated around the bundle axis. See also Fig. 5B-D and accompanying text.

**Supplementary Video 4. Sub-tomogram averages of long and short-pitch F-actin filaments within growth cones.**
3D sub-tomogram averages calculated from volumes of long and short-pitch F-actin and fitted models of F-actin (PDB: 7BT7, (Kumari et al., 2020)) or F-actin-cofilin

(PDB: 3JOS, (Galkin et al., 2011)), as indicated. The video starts with the F-actin model fitted into the long-pitch F-actin sub-tomogram average, showing a good match. The F-actin model is then swapped for the F-actin-cofilin model, that shows a poor match, with no density accounting for cofilin subunits. The F-actin sub-tomogram average is then swapped for the short-pitch F-actin sub-tomogram average, showing a good match, with extra density where cofilin binds. Mesh density is coloured to match within 10 Å of the underlying atomic models.
